# Beyond Static Screens: optical pooled screening of signaling dynamics using time-lapse FLIM

**DOI:** 10.1101/2025.07.11.664338

**Authors:** Sravasti Mukherjee, Menno van Tooren, Dimitris Sfakianakis, Dominique Kemps, Jeffrey Klarenbeek, Giulia Zanetti, Hendrik J. Kuiken, Cor Lieftink, Bram van den Broek, Roderick L. Beijersbergen, Kees Jalink

## Abstract

Genetic screens are powerful tools for linking gene perturbations to cellular phenotypes, yet most pooled imaging approaches rely on static endpoint measurements. Here, we extend optical pooled screening to dynamic signaling phenotypes by combining time-lapse Förster resonance energy transfer–fluorescence lifetime imaging microscopy (FRET-FLIM) with optical enrichment and downstream genotyping.

To enable high-throughput live-cell screening of dynamic phenotypes, we optimized and benchmarked all major steps in the pipeline. We developed FAST-HIPPOS, a Fiji-based automated analysis pipeline that rapidly segments hundreds of thousands of cells, extracts single-cell lifetime time traces, and identifies phenotypic hits in minutes for targeted photoactivation. We further established a robust phototagging strategy using the photoactivatable dye PA-JaneliaFluor646, providing bright, cloning-free labeling across diverse cell types. Benchmarking further demonstrated high sorting fidelity with low false-positive rates and efficient recovery of guide RNAs from as few as 200 sorted cells with minimal PCR bias.

Applying this platform to cAMP signaling in HeLa cells using a custom CRISPR library recovered the expected primary regulators of β-adrenergic signaling—ADRB2, GNAS, and ADCY6—with no gene-level false positives. Together, these optimizations establish a robust general framework for screening dynamic phenotypes that is easily adaptable to other live-cell readouts beyond FRET-FLIM and cAMP.

## INTRODUCTION

Genetic screens link targeted perturbations to measurable phenotypes and have become indispensable for dissecting gene function. Two main formats are commonly used: arrayed and pooled screens. In arrayed screens, a single gene is perturbed in all cells in a well, and each well is analysed separately for a phenotype. While highly informative, these screens are labor-intensive, expensive, and difficult to scale—especially when phenotypic recording is time-consuming. Pooled screens overcome many of these limitations by delivering a library of perturbations to a cell population at low multiplicity of infection (MOI), ensuring that each cell carries ideally a single perturbation^1^. A major advantage of pooled screens is that all cells grow under identical culture conditions, receive the same treatment, and are analysed in parallel. This eliminates variability due to medium evaporation, cell aging, confluence, and day-to-day fluctuations that often hamper arrayed screens^2^.

Pooled screens are used to identify cellular fitness phenotypes, typically through drop-out or enrichment strategies^1^. Identification of genes that affect non-fitness phenotypes often requires imaging-based approaches. Optical pooled screens (OPS)^3^ leverage high-throughput, high-content microscopy techniques to enable unbiased assessment of basically any phenotype discernible by light microscopy, including differentiation, morphology, or protein localization. However, because these screens do not rely on survival, they face the challenge of correctly matching each imaged phenotype to its causative genetic perturbation (guide RNA (gRNA) sequence).

OPS address this challenge in two main ways. One approach involves imaging the phenotypes and then reading the perturbation (gRNA sequence) in each imaged cell via in situ sequencing (ISS)^4–6^. Alternatively, optical enrichment strategies identify cells displaying a phenotype of interest during live imaging and irreversibly tag them with a photoactivatable (PA) or photoconvertible (PC) fluorescent protein. These tagged cells are then isolated by fluorescence-activated cell sorting (FACS), and their genetic perturbations can be identified by next-generation sequencing (NGS)^7–9^.

To date, OPS in mammalian systems have largely focused on endpoint or binary phenotypes such as nuclear translocation, organelle morphology, or protein abundance and localization^7–11^. While highly informative, many biological processes are not defined solely by response magnitude but also by their temporal dynamics. For example, both ERK signaling and neuronal Ca^2+^ signaling demonstrate that sustained signals can produce different cellular outcomes than transient or oscillatory signals^12,13^. Understanding cellular signals, therefore, requires methods capable of capturing the temporal progression of responses in a screening context.

We previously demonstrated that FRET-FLIM imaging can quantify signaling dynamics in an arrayed screening format^14^. We showed that signal amplitude, noise levels, drift, and image analysis speed now suffice for high-throughput single-cell quantification of signaling dynamics. In the present study, we extend the scale of such screens beyond a few dozen genes by optimizing the optical enrichment-based OPS pipeline^7–9^ to detect dynamic phenotypes (Fig. 1A).

**Figure 1.**
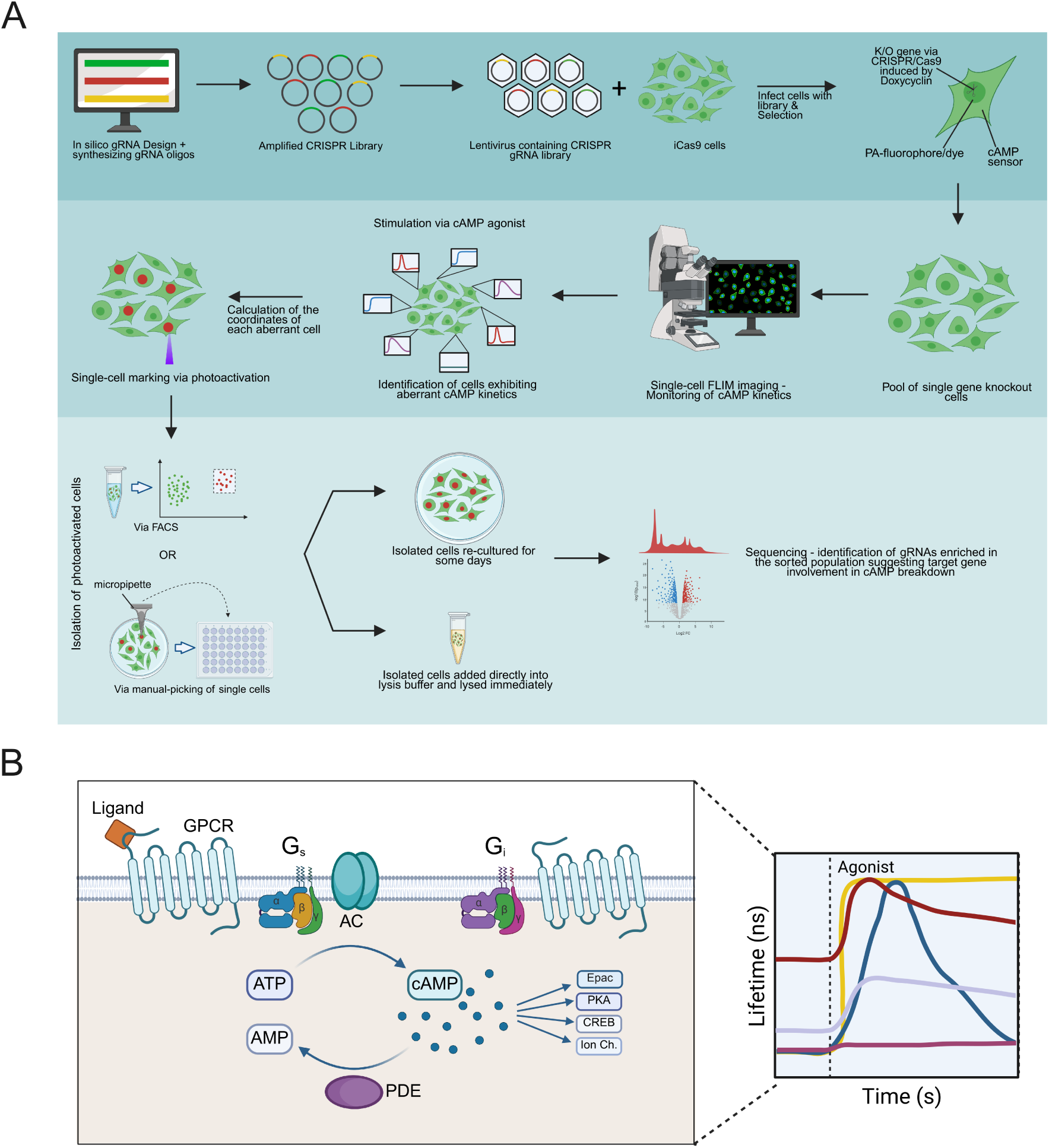
Schematic of the dynamic OPS workflow applied to cAMP signaling. **(A)** OPS workflow. Cells expressing iCas9 (inducible Cas9), the cAMP sensor, and a phototag are transduced with a CRISPR gRNA library and selected. Gene editing is induced with doxycycline, generating a pool of single-gene knockout cells. Cells are screened by time-lapse FLIM, and those showing aberrant cAMP dynamics are identified as hits. Hit cells are revisited and phototagged, followed by isolation via FACS and gRNA identification by NGS. **(B)** Simplified overview of the cAMP signaling pathway. cAMP is generated by Adenylate Cyclases (AC; 10 isozymes) in cells. ACs are activated via the G protein Gα_s_ downstream of specific GPCRs and inhibited by other GPCRs via Gα_i_. The balance of Gα_s_ and Gα_i_ determines cAMP synthesis. cAMP activates several effector proteins, including Protein Kinase A (PKA), cAMP Response Element (CREB), Exchange Protein directly activated by cAMP (Epac), and ion channels. Signal termination is mediated by phosphodiesterases (PDEs, >20 isozymes encoded by 11 gene families^20^. Inset – schematic representation of dynamic cAMP recordings in single cells using the Epac^H250^ FLIM sensor.

Extending OPS to dynamic phenotypes introduces several challenges. Dynamic measurements impose a trade-off between imaging screen throughput and temporal resolution, often limiting the number of cells that can be screened compared to endpoint assays. It also requires analysis software that is fast and robust enough to analyse thousands of single-cell time traces and accurately identify “hit” cells for immediate photoactivation. Finally, existing optical enrichment strategies frequently rely on genetically encoded photoactivatable proteins, which suffer from variable expression levels, loss of plasmid integration^15^, and the need for time-consuming stable cell line generation.

To address these challenges, we implemented several optimizations. First, we replaced genetically encoded phototags with the photoactivatable dye PA-JaneliaFluor646 (PA-JF646)^22^, which provides brighter and less variable labeling, avoids cloning, and can be applied to any cell line within minutes. Second, we developed FAST-HIPPOS, a Fiji-based automated image analysis pipeline that rapidly and reliably identifies hit cells based on user-defined dynamic phenotypes and determines their spatial coordinates for targeted photoactivation. Finally, we optimized the recovery of gRNA sequences from small numbers of sorted cells to ensure NGS accuracy.

Using this improved dynamic OPS pipeline, we performed a screen in HeLa cells expressing our Epac-based FRET-FLIM sensor^16,17^ to measure cAMP signaling dynamics. cAMP is a crucial regulator of diverse cellular functions, including metabolism, gene expression, and differentiation. cAMP signaling is triggered when agonists bind to G-protein coupled receptors (GPCRs) that activate the G protein Gα_s._ Gα_s_ acts as a molecular switch which, in turn, activates Adenylate Cyclase (AC), the enzyme that generates cAMP from adenosine triphosphate (ATP). In this study, we employed isoproterenol to activate Adrenergic β1 and β2 receptors (ADRB1 and ADRB2) present in Cos7 and HeLa cells. Over time, cAMP levels return towards basal levels through the action of phosphodiesterases (PDEs), which hydrolyze cAMP to adenosine monophosphate (AMP) (Fig. 1B). FRET-FLIM imaging provides a unique window into all temporal aspects of cAMP signaling: baseline levels, peak intensities, rise- and fall-times, and more^18^.

Our dynamic OPS platform used a custom CRISPR library targeting 318 genes, most of which are known or predicted components of the cAMP network. Our screen recovered well-known regulators of cAMP^19^ – the Adrenergic Beta Receptor 2 (ADRB2), G protein Gα_s_ (GNAS), and Adenylate Cyclase 6 (ADCY6) - thereby validating the reliability of our platform. While demonstrated here using FRET-FLIM, dynamic OPS is a versatile paradigm adaptable to any dynamic phenotype detectable by microscopy and automated image analysis. This approach therefore provides a general strategy for dissecting gene function in the context of dynamic cellular signaling responses.

## RESULTS

### Cell lines generated for dynamic OPS

First, we generated Cos7 (green monkey kidney) and HeLa (human cervical cancer) cell lines expressing the Edit-R-icas9 plasmid and the Epac-S^H250^ reporter construct^18^. While the screens were carried out with a human gRNA library in HeLa cells, initial technical characterization and optimization of the screening pipeline were carried out in Cos7 cells.

The Epac-S^H250^ plasmid contains: 1) our Epac-S^H189^ high-affinity cAMP sensor, which is dedicated for FLIM readout of FRET signals^17^; 2) a P2A self-cleaving peptide; 3) a nucleus-targeted photoactivatable mCherry (H-NS-PA-mCherry^21^) for cell tagging, and 4) a G418 selection cassette (Fig. 2A). We thoroughly tested sensor FRET span and the precision and throughput of FLIM imaging^18^. The P2A sequence ensures equal transcription of the FRET sensor and the PA-mCherry. However, even though a very bright monoclonal cell line (Cos7^H250^) was selected for further experiments, in our hands, the H-NS tagged PA-mCherry showed poor performance in that its variability increased dramatically over time (Fig. 2B). In addition, fluorescence levels declined markedly well within 3 months of passaging (Fig. 2C), consistent with previously reported instability of integrated fluorescent reporter constructs in long-term culture¹⁵. While this variability does not affect FLIM measurements of cAMP levels because fluorescence lifetime is largely independent of fluorophore concentration, it presents a significant challenge for phototagging. Even minimal accidental photoactivation (with 405 nm UV light) of a bright bystander cell (e.g., when the nuclei of a hit cell and a wild-type bystander are in close apposition) can render the bystander as bright as a fully photoactivated hit cell with much lower expression, thereby compromising the specificity of downstream cell sorting. To overcome these limitations, we replaced genetically encoded phototags with the photoactivatable dye PA-JaneliaFluor646 (PA-JF646)^22^, which we subsequently optimized for use in the dynamic OPS pipeline.

**Figure 2.**
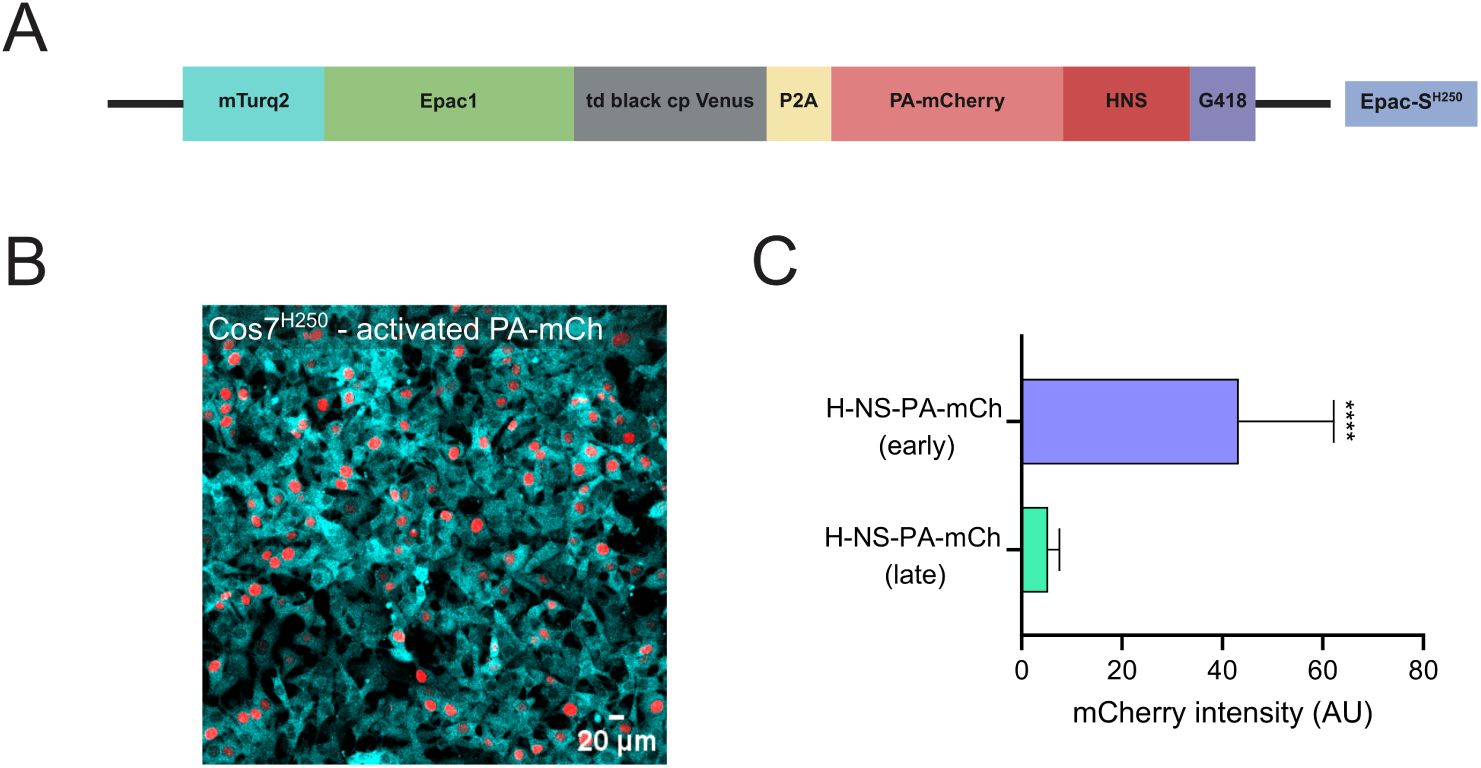
Reporter construct and variability in PA-mCherry expression. **(A)** The reporter construct (Epac-S^H250^) used to express the sensor and a nucleus-targeted (H-NS) PA-mCherry at equal levels. **(B)** Representative image of Cos7^H250^ cells following bulk photoactivation using a benchtop 395 nm LED. mTurquoise intensity is shown in cyan. mCherry intensity shown in red illustrates variability in phototag signal. Scale bar, 20 µm. **(C)** mCherry fluorescence intensity (AU) following photoactivation with the 405 nm laser line in early-passage (just thawed) and late-passage (12 weeks) Cos7^H250^ cells (n = 33 per condition). Intensity is calculated as the difference (I_post-activation_ − I_pre-activation_) and shown as mean ± SD. Statistical significance was assessed using an unpaired one-tailed t-test (****p < 0.0001).

### Optimization of an alternate phototagging strategy: PA-JaneliaFluor646

All dye optimization experiments were performed in Cos7 WT cells. Cells were loaded with 1 µM PA-JF646 (protocol detailed in Materials & Methods (M&M)). This cell-permeable dye binds covalently to primary amines throughout the cells via an N-hydroxysuccinimide (NHS) group, allowing rapid and cloning-free application across cell lines (Fig. 3A). To evaluate its compatibility with the live-cell screening pipeline, we first tested whether PA-JF646 affected cell viability or cAMP signaling. The dye was not cytotoxic (Fig. S1.1A), showed negligible crosstalk with the Epac FRET sensor (Fig. S1.1B), and did not affect either baseline- or agonist-induced cAMP levels (Fig. S1.1C).

**Figure 3.**
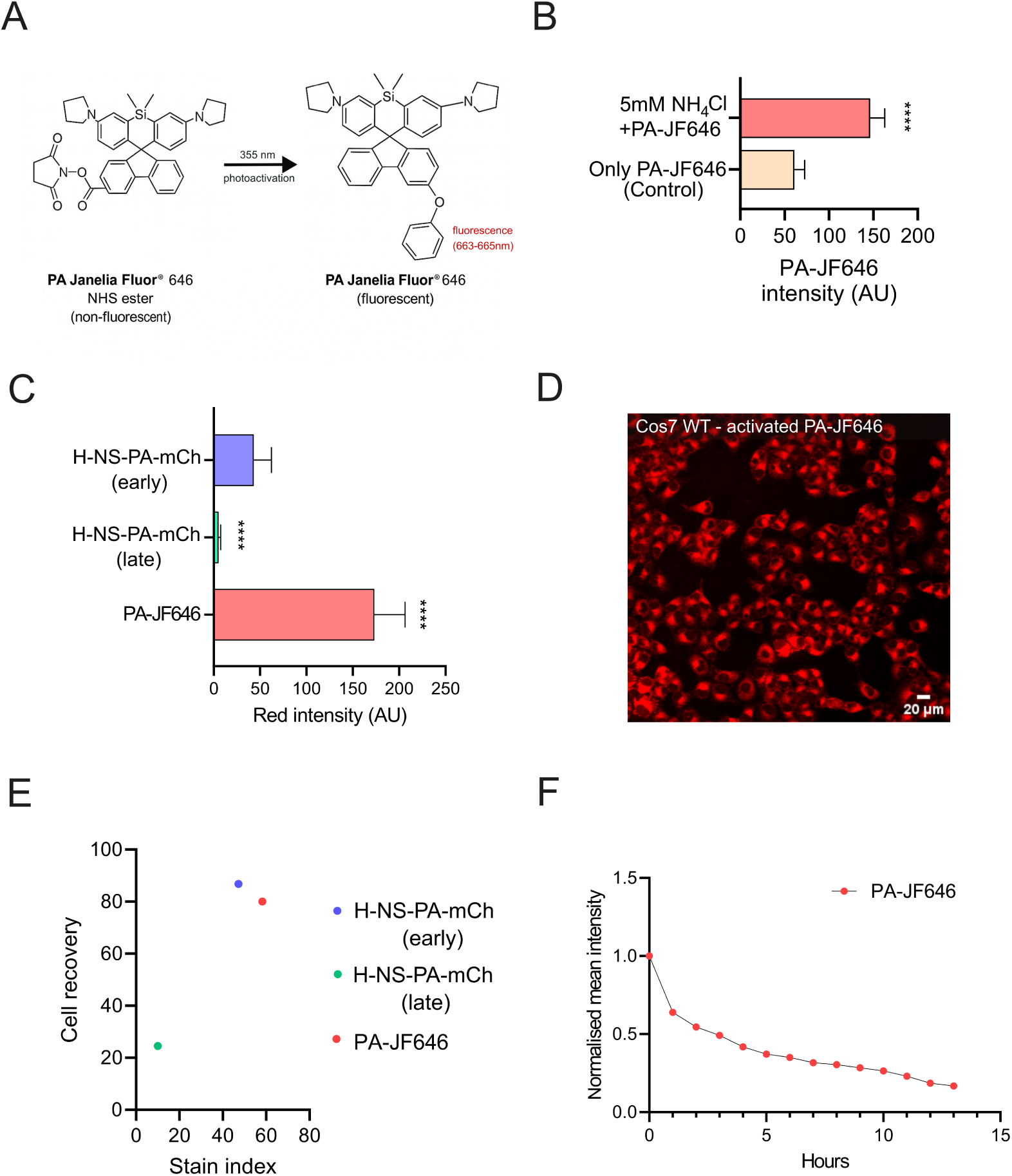
Optimization of PA-JaneliaFluor646 as a phototag. **(A)** Chemical structure of non-activated and activated PA-JF646 dye (taken from Grimm et al., 2016^22^). **(B)** Enhancement of dye labeling by pre-incubation with 5 mM ammonium chloride prior to loading (n ≥ 100 cells). Cells are bulk-activated using a benchtop 395 nm LED. **(C)** Mean red fluorescence (AU) in Cos7^H250^ cells loaded with PA-JF646. For comparison, the data from H-NS-PA-mCherry-expressing early and late cells (Fig. 2C) are also shown. Photoactivation of PA-JF646 was with a 355 nm laser line. Mean ± SD for n = 33 cells. Calculated mean ± SD, CV of the samples: PA-JF646: 173 ± 33, 19%; H-NS-PA-mCherry (early): 44 ± 19, 44%; H-NS-PA-mCherry (late): 5.2 ± 2.2, 43%. **(D)** Cos7 WT cells loaded with PA-JF646 and bulk-activated using a benchtop 395 nm LED, showing homogeneous brightness. Scale bar, 20 µm. **(E)** Correlation between stain index and cell recovery from FACS for different phototagging strategies. Stain index represents the separation (brightness, fold change) between photoactivated and non-activated populations. Cell recovery indicates % of photoactivated cells successfully recovered after FACS sorting (n=2 at least for each condition). **(F)** Signal retention of PA-JF646 in Cos7 WT cells following bulk photoactivation by benchtop 395 nm LED. n > 100. For (B) and (C), Intensity differences (I_post activation_ – I_pre activation_) are shown as mean ± SD; Statistical analysis was with an unpaired, one-tailed t-test. ****p < 0.0001.

Next, photoactivation and dye loading were optimized extensively. Photoconversion of PA-JF646 was with a short pulse of 355 nm laser light. The laser was slightly defocused to ∼ 7 µm spot size so as to cover a significant part of the cell, while not illuminating neighbouring cells (see M&M), resulting in bright fluorescence that appeared confined to individual cells. To optimize dye loading, we took advantage of the fact that NHS reactivity is known to sharply increase at alkaline pH. We therefore devised a loading strategy that transiently elevates intracellular pH via ammonium-driven alkalinization^23^. Accordingly, brief pre-incubation with 5 mM ammonium chloride markedly increased PA-JF646 loading and brightness (Fig. 3B). Cells tagged with PA-JF646 in this manner were substantially brighter than fully activated H-NS-PA-mCherry expressing cells (Fig. 3C; data for HeLa cells in Fig. S1.1D). PA-JF646 signals were also less variable compared to early H-NS-PA-mCherry signals (coefficient of variation (CV)-PA-JF646: 19%; H-NS-PA-mCherry (early): 44%; see legend to Fig. 3C). Reduced variability was also observed during bulk photoactivation (Fig. 3D vs 2B).

FACS experiments showed similar recovery of PA-JF646 cells compared to early-passage H-NS-PA-mCherry cells (Fig. 3E). Additional characterization showed that activated PA-JF646 remained detectable for several hours with approximately 50% of the initial signal retained after four hours (Fig. 3F). Spurious activation of the dye due to ambient light or Epac sensor excitation (Fig. S1.1E) was minimal, and activation of neighbouring cells during phototagging was about 2 orders of magnitude less (Fig. S1.1F).

Together, these results demonstrate that PA-JF646 provides a robust and cloning-free phototagging strategy suitable for OPS experiments.

### FAST-HIPPOS: an automated FLIM analysis and hit detection platform for time-lapse screens

In our previous study^14^, we established an analysis routine using Python and Fiji^24^ to segment single cells and analyse lifetime data. While effective for inspecting individual cell responses in an arrayed format, this approach was too slow for real-time analysis and hit detection in OPS. To address this limitation, we developed FAST-HIPPOS (FLIM Analysis of Single-cell Traces for Hit Identification of Phenotypes in Pooled Optical Screening). FAST-HIPPOS rapidly processes thousands of single-cell time-lapse FLIM traces and identifies candidate hit cells together with their spatial coordinates, enabling targeted photoactivation during the screening workflow. It reads raw Leica (.lif) files, segments cells using CellPose^25^, and extracts single-cell lifetime traces based on intensity-weighted pixel averages. Next, a multi-parameter analysis of the traces can be performed using user-defined criteria such as baseline lifetime, peak response, or average response within a defined time window. Cells fulfilling the specified criteria are classified as hits and visualized through multiple output plots (details in Supplemental Material 2).

To evaluate the performance of the analysis pipeline, we designed a simulation experiment to distinguish a small fraction of signaling-impaired cells from a wild-type (WT) population. First, we generated a polyclonal knock out of ADRB2, the most abundant adrenergic receptor in Cos7 cells, resulting in a heterogeneous pool of biallelic, monoallelic, or in-frame edited cells. Then, we quantitatively compared their cAMP responses following stimulation with 40 nM isoproterenol with those of Cos7^H250^ WT cells (Fig. 4A). As expected, WT cells exhibited a homogenous, sustained cAMP response that saturated the FRET sensor, whereas a large fraction of ADRB2 polyclonal KO cells showed significantly reduced responses. Crucially, these KO cells retained normal forskolin-induced activation, confirming that the downstream adenylyl cyclase machinery remained functional. To discriminate between the two cell populations, we selected a threshold of 3.05 ns in the mean response lifetime (measured over the 12 minutes preceding the calibration, i.e., frames 12–24). With this criterion, only 1.8% of WT cells were categorized as hits (Fig. 4A, left-top panel), whereas 41.1% of the ADRB2 polyclonal KO population met the hit criteria, reflecting the phenotypic penetrance in the polyclonal KO population (Fig. 4A, right-bottom panel).

**Figure 4.**
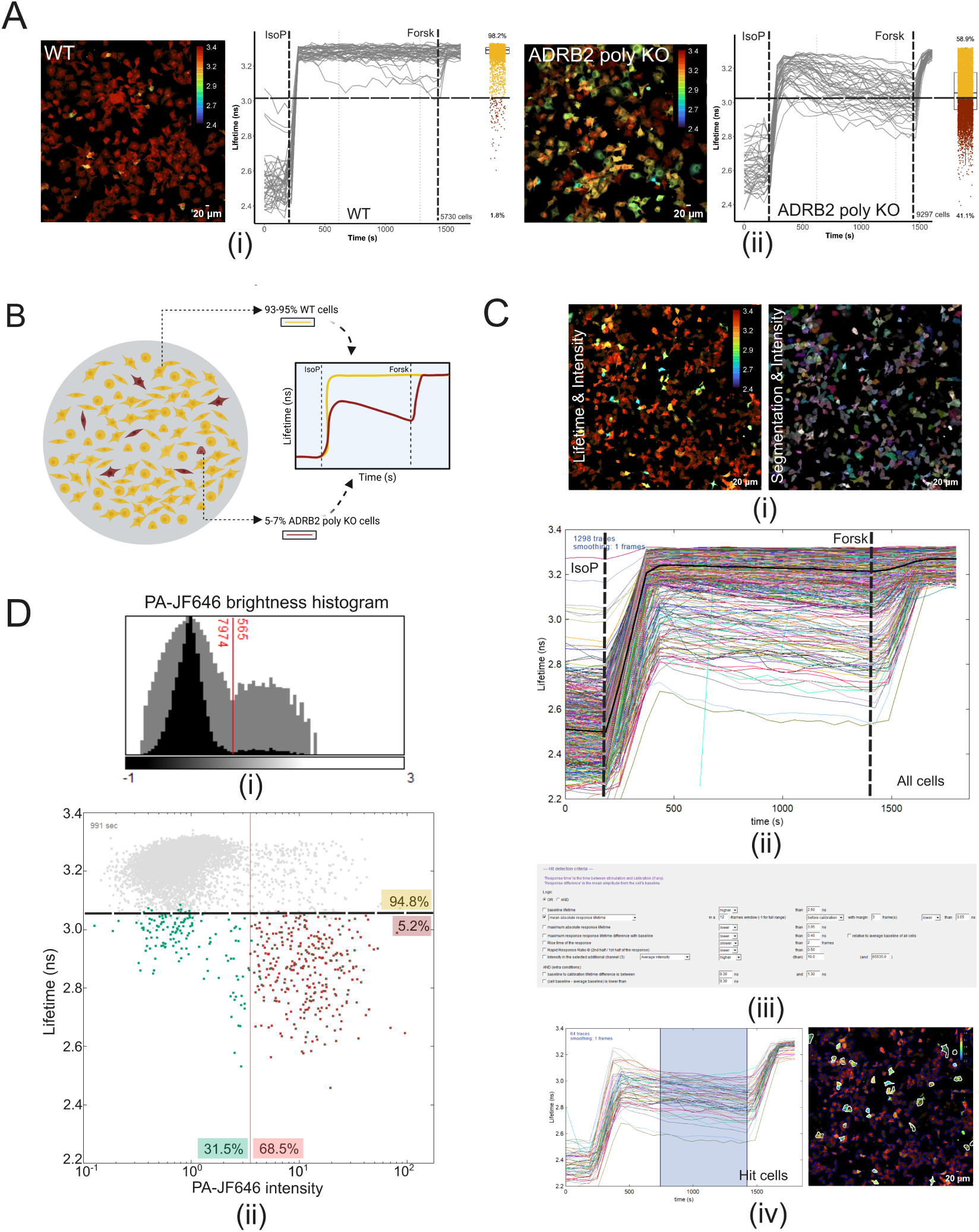
FAST-HIPPOS performance in a simulated pooled screening experiment. **(A) (i), (ii)** Response of Cos7^H250^ WT and Cos7^H250^ ADRB2 polyclonal KO cells after stimulation with 40 nM isoproterenol and calibration with 25 µM forskolin as indicated. Shown are 45 randomly-selected representative FLIM traces. The boxplots show the lifetimes of all imaged individual cells averaged over frames 12-24 (∼12 minutes; grey dotted lines) just before calibration. Quantification: WT, 1.8% of cells have a mean lifetime < 3.05 ns; ADRB2 polyclonal KO cells, 41.1 % of cells have a lifetime < 3.05 ns. Also shown are representative FLIM images taken just before calibration. Turbo LUT used for FLIM image ranges from low lifetime (2.4 ns; blue) – high lifetime (3.4 ns; red); Scale bar, 20 µm. **(B)** Schematic representation of experimental design. ∼7% PA-JF646-labelled ADRB2 polyclonal KO cells are mixed with unlabeled WT cells to simulate rare hits in a pooled screen. **(C) (i)** Lifetime image of mixed cell population (left) and segmentation overlay (right). Scale bar, 20 µm. **(ii)** Lifetime time traces of a subset (1/8^th^ of the mosaic image; 1298 segmented cells) showing stimulation with 40 nM isoproterenol and calibration with 25 µM forskolin. The timepoints of stimulation and calibration are automatically calculated from the average of all traces (thick black dotted line) or can be entered manually. **(iii)** Hit selection dialog showing various selection criteria, which can be logically combined to enable screening for a large variety of dynamic phenotypes. **(iv)** Visualization of identified hit cells and corresponding lifetime traces. The frames used for analysis are highlighted in blue. **(D) (i)** Histogram of PA-JF646 intensity (X-axis plotted in log(10) scale) used for automatic Otsu thresholding. **(ii)** Scatter plot of lifetime versus PA-JF646 intensity at frame 17 (randomly chosen) showing true positives. i.e., labeled KO cells (red, 68.5%) and false positives (green, 31.5%). Hit selection is based on mean response lifetime (frames 12-23) < 3.05 ns. A total of 5.2% cells were detected as hits. IsoP, 1 µM isoproterenol; Forsk, 25 µM forskolin. The total number of analysed cells is indicated in each time-trace graph.

Finally, for the simulation experiment, the ADRB2 polyclonal KO cells were pre-labeled with photoactivated PA-JF646 to distinguish them from the WT cells. After trypsinization, ∼7% of ADRB2 polyclonal KO cells were seeded onto sub-confluent WT cells in glass-bottom dishes to simulate rare hits in a pooled screening experiment (Fig. 4B). The mixed population was imaged for 30 minutes, stimulated with 40 nM isoproterenol, and calibrated with 25 µM forskolin. Segmentation (Fig. 4C (i)), extraction of lifetime traces (Fig. 4C (ii)), and hit detection using the above-defined criterion (3.05 ns) by FAST-HIPPOS (Fig. 4C (iii), (iv)) identified 5.2% of the total population as hits. To verify, Otsu thresholding^26,27^ of red fluorescence intensity showed that 68.5% of these detected hits were true positives (PA-JF646-labeled KO cells), demonstrating substantial detection accuracy (Fig. 4D (i), (ii)). The identification of 5.2% hits is in reasonable agreement with our experimental design (for details, see M&M).

Note that FAST-HIPPOS ranks hits according to phenotypic strength, allowing prioritization of the most robust candidates within the ∼1000-cell photoactivation limit imposed by imaging time constraints. This effectively tightens the threshold if more than 1000 hits are detected. In addition, false positives from accidental bystander activation are rejected by stringent gating during FACS prior to sequencing, ensuring reliable downstream identification of enriched perturbations. Importantly, a detection accuracy of ∼68 % proved sufficient for the screening workflow.

### Estimating FACS false-positive rates and NGS sensitivity: a Dolcetto library benchmark

One of the main trade-offs in dynamic OPS is that only a limited number of cells can be imaged at sufficient temporal resolution (see Supplemental Material 3). Consequently, the number of photoactivated hit cells recovered from a screen is often relatively small, which places strict requirements on genomic DNA recovery and downstream sequencing. Inefficient DNA extraction can lead to loss of true hit gRNAs, whereas low DNA input may introduce PCR amplification biases or random overrepresentation of specific guides.

To determine the sensitivity of gRNA detection and the specificity of cell sorting, we used the Dolcetto gRNA library^28^. The Dolcetto library contains two independent gRNA sets: Set A (57,050 gRNAs) and Set B (57,011 gRNAs), with no overlapping guides. Cos7^H250^ cells were transduced separately with lentiviruses encoding either Set A or Set B at a MOI of approximately 0.3 and 0.4, respectively, ensuring that most of the transduced cells received only a single gRNA. Following puromycin selection, the photolabels in cells transduced with set A were activated with a benchtop 395 nm LED and mixed with non-activated set B cells at a 1:4 ratio (set A: set B). Different numbers of photoactivated cells were isolated by FACS (200, 500, 2000, 5000, and 10,000 per sample) in triplicate (Fig. 5A). Following sorting, genomic DNA (gDNA) was isolated from these cells by using a DirectPCR cell lysis reagent (as opposed to kit-based gDNA isolation which results in significant DNA loss) and a 2-step PCR was performed using a low-fidelity, crude lysate compatible Taq polymerase (as opposed to high-fidelity Phusion polymerases), to amplify the target sequence and add adapters for NGS (detailed protocol in M&M).

**Figure 5.**
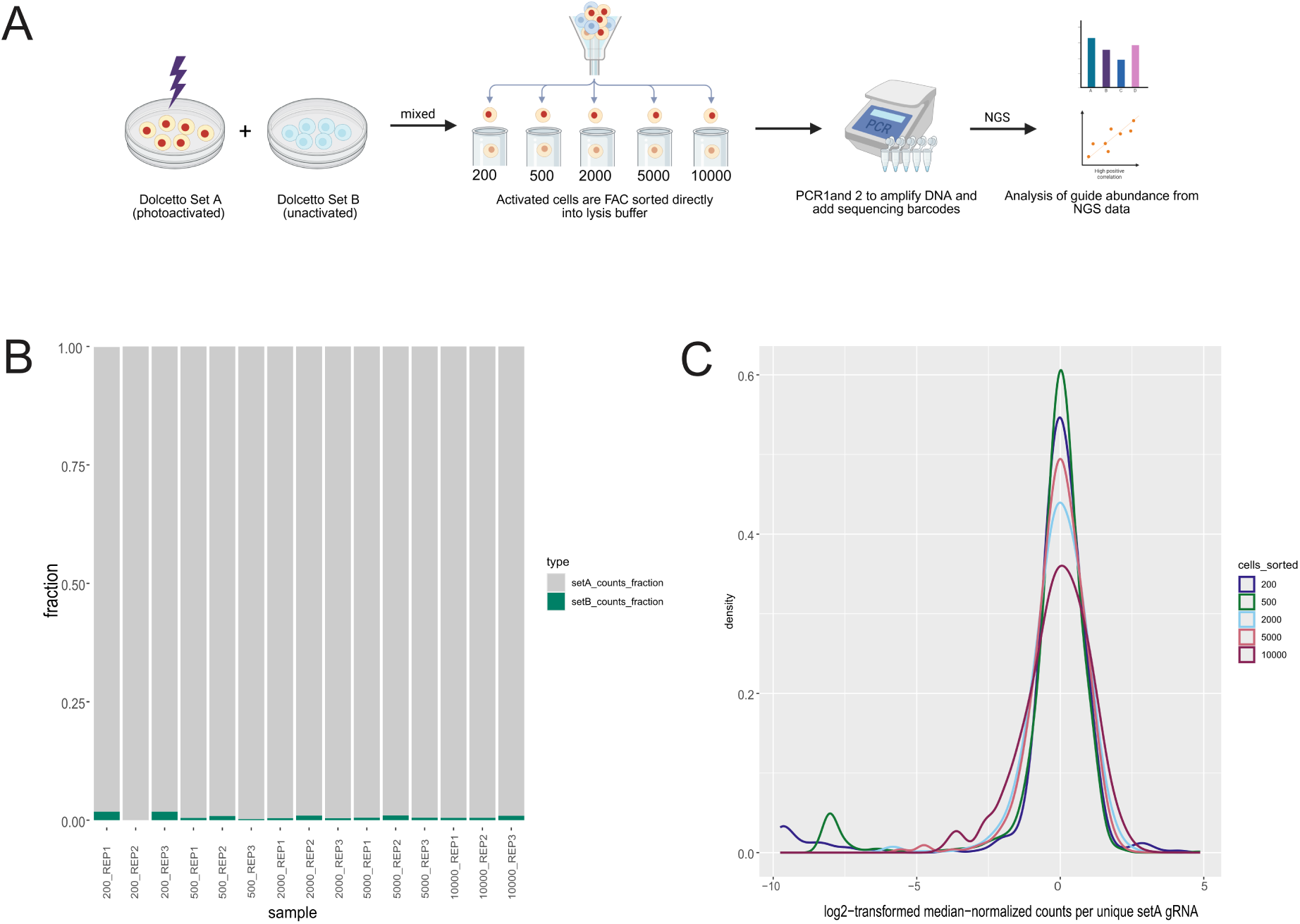
Dolcetto library benchmark for gRNA recovery and sorting specificity. **(A)** Schematic representation of experimental flow. Photoactivated Dolcetto Set A cells are mixed with non-activated Set B cells, followed by FACS isolation of varying numbers of activated cells (200–10,000). Sorted cells are lysed, and gRNAs are amplified and sequenced. **(B)** Fraction of sequencing reads mapping to Set A and Set B gRNAs, demonstrating low contamination from non-activated cells. **(C)** Normalized count distribution of individual gRNAs from Set A following sorting and sequencing, indicating minimal amplification bias. Representative example from triplicate experiments.

The NGS results showed that most of the sequence reads (98.2 – 100%) were unique gRNA sequences and mapped to gRNAs of set A (97.4 – 100%). This indicates that contamination from Set B guides was minimal, corresponding to an estimated false-positive rate below 2.6% in the worst case, and typically below 1% (Fig. 5B). Normalized gRNA count distributions (Fig. 5C) showed no obvious overrepresentation of individual gRNA sequences, indicating that the amplification protocol did not introduce strong PCR biases despite the relatively high number of amplification cycles. In addition, control experiments confirmed that the use of low-fidelity Taq polymerase did not introduce detectable sequence errors in the amplified gRNAs (Fig. S1.2).

In this experiment, the gRNA sequences served purely as DNA barcodes, and therefore, it was not relevant that the Dolcetto library targets human genes while the host cells were Cos7. Because the experiments used early-passage Cos7^H250^ cells expressing H-NS-PA-mCherry, phototagging was performed using PA-mCherry rather than PA-JF646.

Together, these experiments demonstrate that the dynamic OPS workflow enables reliable recovery of gRNA sequences from small numbers of sorted cells while maintaining a low false-positive rate during cell sorting.

### Pilot screen for elevated baseline fluorescence lifetime in HeLa^H250^ cells

To validate the performance of the optimized screening pipeline, we performed a pilot screen using a moderate-sized custom-made cAMP gRNA library targeting 318 genes with known or expected roles in shaping the dynamics of receptor-mediated cAMP signals (318 genes, 1272 gRNAs; details of library design are in Supplemental Material 4). As a test phenotype, we screened for genes that increase fluorescence lifetime in unstimulated HeLa^H250^ cells (i.e., baseline lifetime). We deliberately chose a phenotype for which the library contains a strong positive control – Rap guanine nucleotide exchange factor 3 (RAPGEF3/Epac1), the cAMP-binding domain in our FRET-FLIM sensor. The targeting gRNAs are predicted to remove most of the RAPGEF3/Epac1 moiety from our sensor, together with the fluorescent acceptor of the probe (Fig. S1.3). As a result, cells retain only the mTurquoise2 donor fused to a truncated Epac fragment. In the absence of FRET, the donor exhibits a high fluorescence lifetime (baseline), providing a robust and unambiguous screening phenotype.

Cells were prepared for the screen as described in M&M. Baseline fluorescence lifetime of unstimulated HeLa^H250^ cells was recorded over two imaging frames covering 304 tiles per dish (∼42,000 cells), requiring approximately 12 minutes of imaging time. For each of 3 biological replicates, data from 4 dishes were pooled and automatically analysed (see M&M, Table 3), achieving ∼140x coverage of each gRNA per replicate. In addition, a control dish was stimulated with forskolin in each replicate to verify cell health and confirm the dynamic range of the FRET sensor; thus, a total of 15 dishes were imaged.

Automated analysis identified an average of 170,000 cells per biological replicate. Hits were defined as cells displaying a baseline lifetime differing from the population average (2.27 ns) by more than 4 standard deviations (S.D. = 0.077), corresponding to a threshold of τ_b_ > 2.6 ns (Fig. 6A). Cells fulfilling this single criterion were photoactivated, isolated by FACS, and subjected to NGS analysis. The primary hit selection was based on a statistical analysis with DESeq2 and MAGeCK’s Robust Rank Algorithm. RAPGEF3/Epac1 was identified as the primary hit (Fig. 6B), thereby confirming the robustness of our screening pipeline.

**Figure 6.**
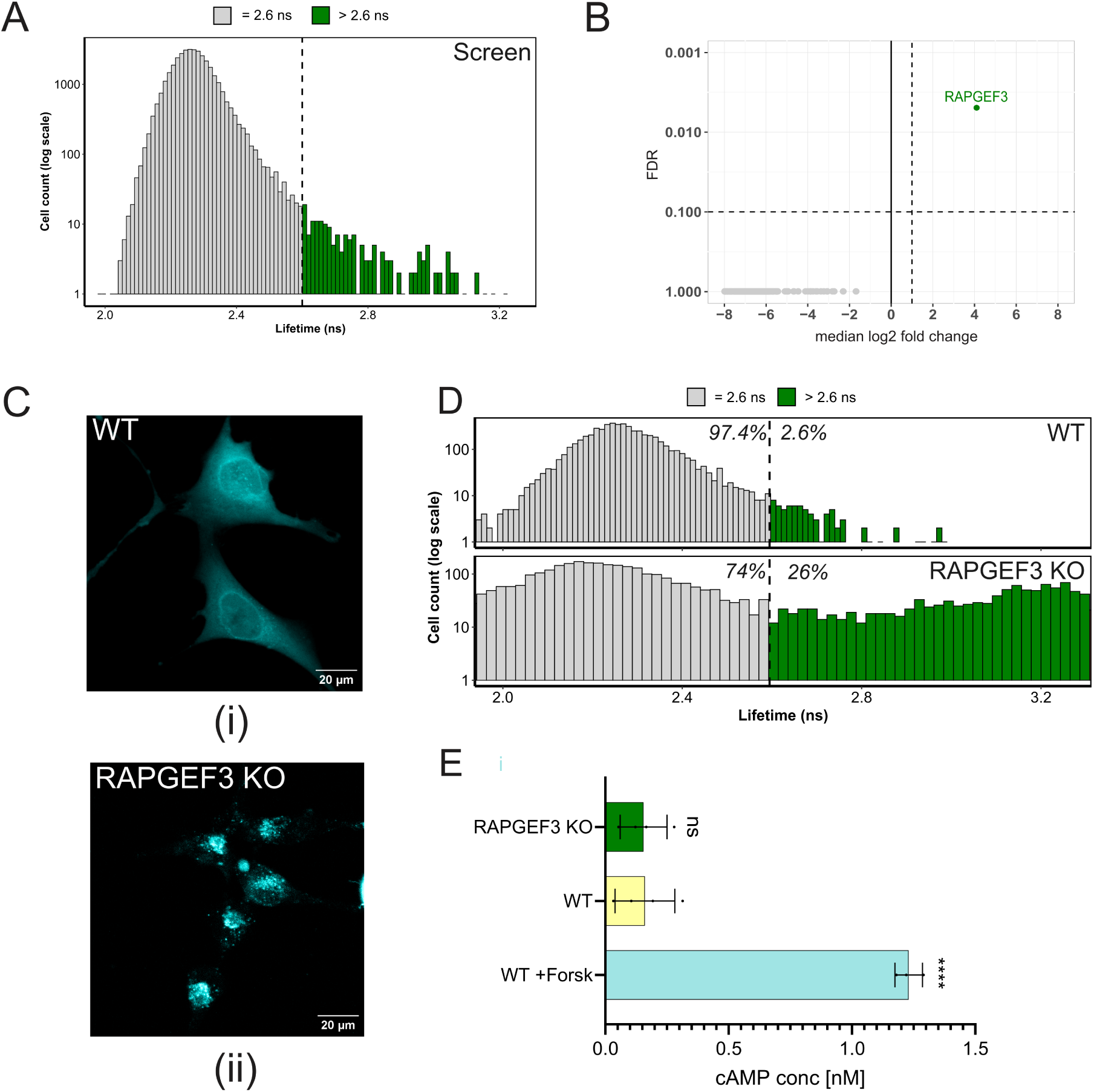
Pilot screen for increased baseline lifetime and validation of RAPGEF3/Epac1. **(A)** Example distribution of baseline lifetimes (in dish 1 of experimental replicate 1) of HeLa^H250^ cells transduced with the cAMP library. A threshold of 2.6 ns (green) was used to define hits. **(B)** Gene enrichment analysis identifying RAPGEF3/Epac1 as the primary hit based on gRNA enrichment (FDR ≤ 0.1; median log₂ fold change ≥ 1). **(C) (i)** HeLa^H250^ WT (control) cells expressing the Epac sensor and **(ii)** HeLa^H250^ RAPGEF3/Epac1 polyclonal KO cells with erroneous sensor localization. mTurquoise intensity is depicted in cyan. Scale bar, 20 µm. **(D)** Distribution of baseline lifetimes in HeLa^H250^ WT (top) and HeLa^H250^ RAPGEF3/Epac1 polyclonal KO cells (bottom). Approximately 2.6% of WT cells and 26% of KO cells exceed the 2.6 ns threshold. Representative experiment from n = 4 independent experiments. **(E)** ELISA measurement of basal cAMP levels in HeLa^H250^ WT and RAPGEF3/Epac1 polyclonal KO cells. Forskolin-treated WT cells serve as a positive control. Data are shown as mean ± SD (n = 3). Statistical significance was assessed using an unpaired one-tailed t-test (ns, not significant; ****p < 0.0001).

#### Validation of the hit

For validation, a HeLa^H250^ RAPGEF3/Epac1 polyclonal KO cell line was generated, confirmed by Sanger sequencing, and tested in FLIM experiments and a cAMP ELISA assay. The KO cells showed aberrant fluorescence distribution compared to WT cells (Fig. 6C (i), (ii)). FLIM analysis of RAPGEF3/Epac1 polyclonal KO cells showed very high baseline lifetime values (∼26% of cells showed lifetimes > 2.6 ns) compared to ∼2.6 % in WT cells (Fig. 6D). In line with this notion, the ELISA analysis indicated normal basal cAMP levels in RAPGEF3/Epac1 polyclonal KO cells (Fig. 6E), proving that the high lifetime values are due to the unquenched donor lifetime (mTurquoise2) and not due to increased cAMP levels.

Having established that the screening pipeline reliably detects known perturbations, we next applied the workflow to identify regulators of agonist-induced cAMP signaling dynamics.

### Dynamic OPS identifies genetic regulators of cAMP response amplitude and duration in HeLa^H250^ cells

Finally, we applied dynamic OPS in HeLa^H250^ cells using our custom cAMP gRNA library to identify genes required for a full and sustained cAMP increase in response to the β-adrenergic agonist isoproterenol.

We first characterized the isoproterenol response dynamics in control (HeLa^H250^ WT) cells to establish wildtype response characteristics (Fig. 7A). Addition of isoproterenol triggered a rapid increase in cAMP after 10 ± 5 s. cAMP then remained high for an average of 15 ± 5 min before gradually returning towards baseline. The time-course of this response also dictates the sampling speed necessary to capture the dynamics, which in turn determines how many cells/fields of view can be included in the screen, given the optimized imaging speed as discussed in Supplemental Material 3.

**Figure 7.**
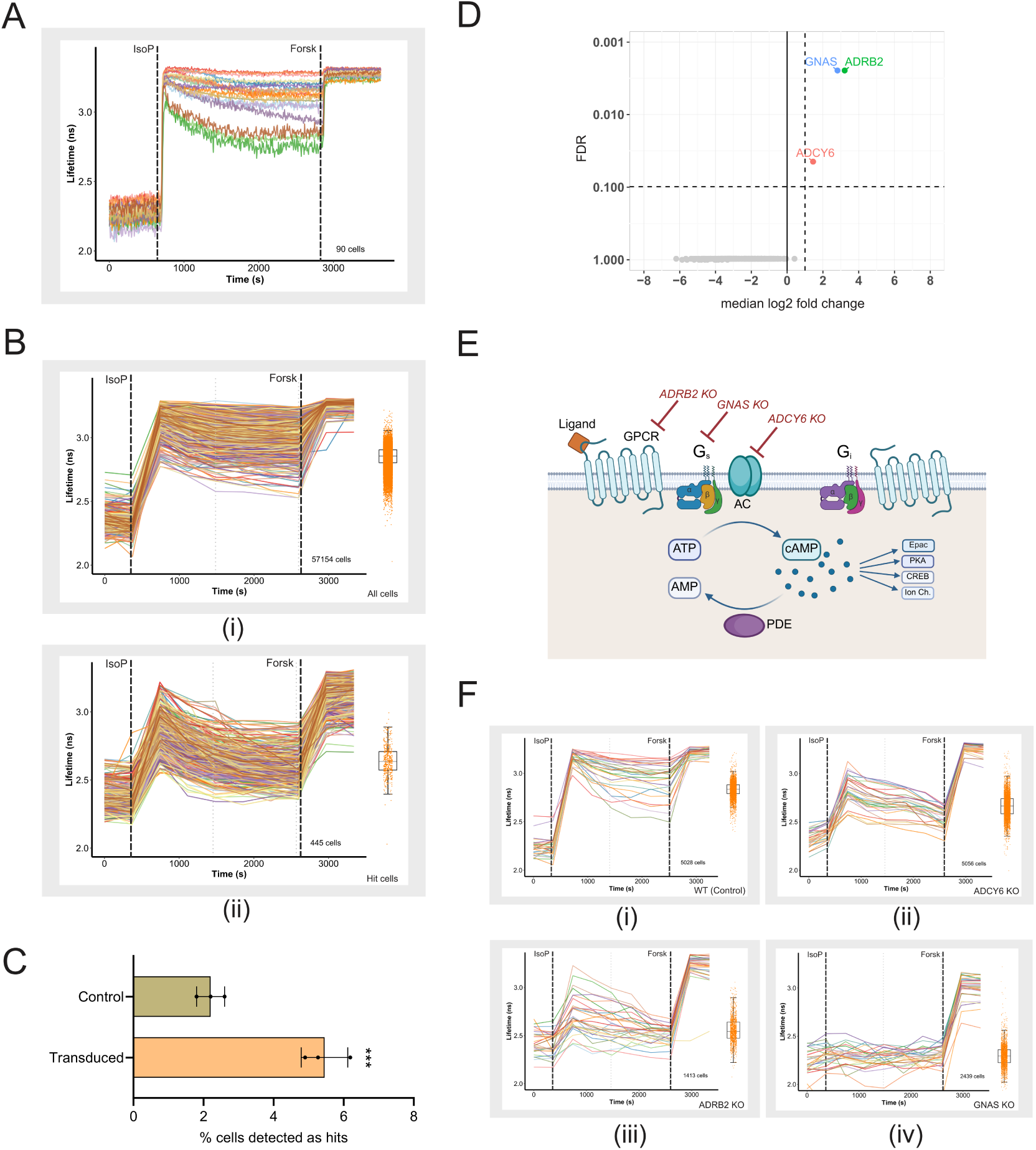
Identification and validation of hits in a dynamic OPS for low cAMP responders. **(A)** Representative FLIM traces (n=30, randomly selected) from HeLa^H250^ WT cells stimulated with 1 µM isoproterenol and calibrated with 25 µM forskolin, imaged at high temporal resolution. **(B) (i)** Representative randomly selected traces and summary boxplot (∼58,000 cells) measured in dish 1 of Replicate 1. **(ii)** Cells identified as low responders in this dish using screening criteria defined in the text. **(C)** Fraction of cells (mean ± SD) classified as hits across 3 transduced biological replicates, compared to WT controls. Statistical significance was assessed with an unpaired, one-tailed t-test; ***p = 0.001. **(D)** Genes significantly enriched with FDR ≤ 0.1; median log₂ fold change ≥ 1 – ADRB2, GNAS, and ADCY6. **(E)** Schematic of the cAMP signaling cascade highlighting the key hits from the screen. **(F)** Representative lifetime traces (n = 30, randomly selected) from validation experiments in WT and KO cell lines: **(i)** HeLa^H250^ WT (Control) (n=4), **(ii)** HeLa^H250^ ADCY6 KO (n=4), **(iii)** HeLa^H250^ ADRB2 KO (n=3), and **(iv)** HeLa^H250^ GNAS KO (n=3). In (B) and (F), boxplots quantify the lifetime of all individual cells, averaged over frames 6-8 (grey dotted lines). IsoP, addition of 1 µM isoproterenol; Forsk, addition of 25 µM forskolin.

Expected hits from this screen would be genes that, in WT cells, positively contribute to either the magnitude or the longevity of cAMP formation, or genes that strongly downregulate cAMP clearance. Thus, knockout of these genes should result in low-amplitude isoproterenol responses or short-duration transient responses. A 6-minute interval suffices to capture the relatively slow sustained phase of the response (Fig. 7B (i)) and allows a total of 240 imaging tiles (∼30,000 cells) to be acquired. An hour-long time-lapse thus contains 10 frames: baseline (frames 1-2), response (frames 3-8), and calibration (frames 9-10).

For analysis, the following filtering criteria were applied: (i) Cells were selected to occupy at least 220 pixels, causing CellPose to segment on average of 30,000 cells in each of 4 dishes, or ∼120,000 cells per replicate; (ii) cells exhibiting a calibration response (lifetime after forskolin addition minus baseline value) less than 0.3 ns were excluded to reject dead cells and debris; (iii) cells with baseline lifetimes deviating more than 0.3 ns from the population average were excluded from analysis.

Finally, cells were classified as low responders if the mean lifetime during frames 6–8 remained within 0.4 ns of baseline (frames 1-2), corresponding to a reduction of more than 50% compared with the average population response in controls (Fig. 7B(ii)).

With these criteria, the hit detection rate in control samples was ∼2.2%, whereas significantly higher hit rates were detected in the KO samples across three replicates: 5.3%, 4.9%, and 6.2%, respectively (Fig. 7C). This corresponds to an approximate false-positive background of 40%, which is expected given the intentionally permissive detection threshold and the biological heterogeneity and day-to-day variability present in the samples. Hit cells detected in the samples were phototagged, sorted, and sequenced. The primary hit selection was based on statistical analyses using DESeq2 and MAGeCK’s Robust Rank Algorithm. Three genes stood out in HeLa cells: the expected Adrenergic Beta Receptor 2 (ADRB2), G protein Gα_s_ (GNAS), and Adenylate Cyclase 6 (ADCY6) (Fig. 7D). These genes form the core signaling axis of β-adrenergic cAMP signaling (Fig. 7E).

#### Validation of hits

To validate these hits, monoclonal KO cell lines were generated in HeLa^H250^ and confirmed by Sanger sequencing. FLIM measurements showed that ADRB2 and ADCY6 KO cells exhibited strongly reduced cAMP responses, whereas GNAS KO cells showed a near-complete loss of response compared with wild-type controls (Fig. 7F(i–iv)).

HeLa cells express several adenylyl cyclase isozymes, including ADCY3, ADCY6, ADCY7, ADCY9, and ADCY10^29,30^, with partial functional overlap. Our findings suggest that ADCY6 is the dominant adenylyl cyclase coupling to Gα_s_ in HeLa cells (Fig. 7F (ii)). However, ADCY6 KO cells still responded strongly to forskolin, indicating that other ACs remain functional. The partial response in ADRB2 KO cells (Fig. 7F (iii)) was to be expected: isoproterenol is an agonist to both adrenergic receptors, ADRB2 and ADRB1, and thus the remaining response might be mediated by low-level ADRB1 expression, although the Human Protein Atlas reports low expression in HeLa cells^31^. Similarly, the near-complete loss of response in GNAS KO cells (Fig. 7F (iv)) is no surprise because GNAS encodes the only Gα_s_ isoform.

Together, these experiments show that the dynamic OPS workflow successfully identifies key regulators of receptor-mediated cAMP signaling.

## DISCUSSION

In this study, we extend optical pooled screening (OPS) to dynamic cellular phenotypes by integrating time-lapse FRET-FLIM imaging with optical enrichment and pooled CRISPR perturbations. Using this approach, we screened ∼130,000 cells across three biological replicates and achieved ∼100× library coverage within three days. By moving beyond static endpoint measurements^8,32^, the platform enables the identification of genetic regulators that influence specific temporal features of signaling responses. Several stages of the workflow were systematically optimized to achieve this performance.

First, we optimized phototagging of hit cells. Previous OPS studies have typically relied on genetically encoded photoactivatable fluorescent proteins such as PA-mCherry or Dendra2^7,8^. In our implementation, however, PA-mCherry expression proved too variable to serve as a reliable phototag. We therefore replaced it with the photoactivatable dye PA-JF646. This dye is substantially brighter than PA-mCherry and eliminates the need for cloning or stable cell line generation. A limitation is that the activated signal decays faster than protein-based tags^33^, losing ∼50% intensity within four hours, which requires that FACS sorting be performed shortly after screening. For best results, it is recommended to perform the experiments in the dark and shield the samples from direct exposure to ambient light, even though spurious activation is minimal (Fig. S1D).

Second, we developed the software for rapid analysis of dynamic phenotypes. FAST-HIPPOS is a Fiji-based analysis pipeline that segments cells, extracts single-cell FLIM traces, and identifies candidate hit cells within minutes. The software processes ∼30,000 cells from an hour-long time-lapse experiment in ∼eight minutes using GPU-accelerated routines^34^. A detailed description of the analysis pipeline and the script is available at https://imagej.net/plugins/fast-hippos. While developed for automating the analysis of FLIM screens, the analysis framework is readily adaptable to other time-resolved imaging readouts, including intensity- or ratio-based reporters. It can also be easily adapted for analysis of other quantitative dynamic phenotypes discernible by microscopy.

Finally, we optimized genomic DNA recovery. OPS typically yields very limited number of hit cells, and this is worsened by additional losses during FACS, due to sample preparation procedures, stringent singlet gating in the sorter, and to the use of high-purity FACS sorting settings that prioritize accuracy over yield. Owing to FACS losses, other groups performing OPS have adopted alternative strategies for hit isolation, including Photostick^35^ for photochemical cell tethering, single-cell magneto-optical capture (scMOCa) for magnetic-optical capture, and Raft-Seq for robotic cell picking of microraft arrays^3,11,36,37^. Nonetheless, we opted for FACS-based recovery as it was the easiest to set up, and rather focused on improving the extraction and amplification of DNA from low cell numbers. This also broadens the applicability of the technique for screening of rare phenotypes or rare cell populations, such as circulating tumor cells, sorted immune subsets, or developing organoids. Our improvements also present a cost-effective alternative to whole-genome sequencing for screening applications.

The performance of the screening pipeline was demonstrated through a pilot screen for cells with high basal fluorescence lifetimes and a dynamic OPS for low cAMP responders. The predicted strong positive control, RAPGEF3/Epac1, was identified as a hit in the pilot screen, and the entire main axis of the adrenergic receptor response cascade, namely ADRB2, GNAS, and ADCY6, were identified as hits in the dynamic OPS, thereby validating the reliability and specificity of our screening pipeline.

A defining strength of this pooled approach is that all cells transduced with the library are treated and imaged under identical conditions, minimizing experimental variability. By recording at 6-minute intervals with a 20x objective, we achieved high-resolution time-lapse data for ∼30,000 cells per run. This temporal depth enables precise cell segmentation and the application of time-smoothing filters to reduce stochastic noise both for segmentation and for dynamic measurements. This is a significant advantage over static endpoint imaging, where transient artifacts such as floating debris could ruin individual cell data. Furthermore, the screen’s dynamic nature enables stringent built-in quality control. For instance, we required a minimum calibration response (a >0.3 ns shift from baseline values after forskolin addition) to systematically exclude dead cells and non-responsive debris. At the single-cell level, this workflow operates with a relatively high background (∼40% of detected hits correspond to wild-type cells), determined by both the deliberately permissive hit-selection criteria and cellular heterogeneity. However, our workflow prioritizes accuracy by ranking hits based on phenotypic strength and restricting photoactivation to the top 1,000 candidates only. Combined with a robust 100-fold coverage, three biological replicates, strict FACS gating and high-purity sorting, the screen design ensures that such stochastic false positive cells do not propagate to the gene level. Even with minor fluctuations in guide enrichment across replicates for hits like ADCY6, the final hit list showed no gene-level false positives, demonstrating that the workflow is robust to cellular noise and reliably prioritizes genuine regulators. In summary, this study establishes a practical framework for integrating OPS with single-cell imaging of signaling dynamics. While developed here for cAMP signaling using FRET-FLIM, the approach is readily generalizable to any live-cell readout compatible with automated imaging and analysis. By enabling pooled genetic interrogation of temporal signaling features, this platform opens the door to systematic dissection of signaling kinetics in mammalian cells.

## LIMITATIONS OF THE STUDY

This study was not designed to provide an exhaustive analysis of genes involved in shaping cAMP dynamics. Given that the gRNA library was largely composed of genes involved in the cAMP pathway and receptor recycling, with only a limited number of genes lacking known links to cAMP signaling, we expected to recover mainly known pathway components, limiting the potential for discovery of novel targets. Furthermore, hit selection was restricted to the top ∼1000 per sample. While this helps minimize false positives, it likely reduces sensitivity for genes producing subtle or intermediate phenotypes.

The use of strong agonists, such as isoproterenol, leads to a massive accumulation of cAMP and saturates FRET, potentially masking certain aspects of cAMP dynamics. This limitation could be addressed by using FRET-FLIM sensors with a broader range of affinities^38,39^. In addition, the cytosolic FRET-FLIM sensor used here cannot resolve spatially restricted cAMP signals, such as nanodomains or localized hotspots of cAMP turnover.

From a technical perspective, there is ample room for improvement and further development. Photoactivation may be accelerated by activating hit cells in parallel, rather than sequentially, using e.g. a digital micromirror device^8^. A key limitation remains imaging speed: capturing the rapid activation phase of cAMP signaling (∼15 s) would require frame intervals of 5 s or less, which would drastically reduce spatial coverage (∼5 fields of view, ∼2000 cells per assay using a 20x objective). At present, FLIM acquisition speed is constrained by TCSPC-based detection, and it remains unclear whether substantial gains can be achieved without compromising accuracy on a confocal setup.

While FAST-HIPPOS already benefits from GPU acceleration, further gains could be achieved through additional parallelization. Versatility would benefit from inclusion of cell tracking to support longer time-lapses or migratory phenotypes.

Finally, more broadly, the framework presented here can be adapted to screen for diverse dynamic behaviors, including oscillatory signaling, sustained activation, recovery from receptor desensitization, or dynamic changes in cell morphology during e.g. cytokinesis and locomotion.

## MATERIALS AND METHODS

### Making stable Cos7^H250^ and HeLa^H250^ cell lines

To generate stable iCas9-expressing lines, HeLa (#CCL2) and Cos7 (#CRL-1651) cells were cultured in DMEM (+Glutamax) (Gibco, Thermofisher, #61965059) supplemented with 10% FCS. Then, they were transduced with the Edit-R iCas9 lentiviral vector (Dharmacon, #NC1606271) and selected with 10 µg/mL Blasticidin (InvivoGen, #ant-bl-05) for three days to enrich for transduced populations. For generation and verification of the iCas9 cell lines, the protocol described in Neto et al., 2023^40^ is used.

For the creation of stable lines with the Epac sensor, the cells were transfected with the Epac-S^H250^ construct (Addgene, #245739), which contains the Epac-S^H189^ FRET sensor and H-NS-PA-mCherry^18,21^, separated by a self-cleaving 2A sequence^41^. This construct was transfected using the Tol2 transposon system^16,42^. Transfection was with 2 plasmids: one containing a cDNA carrying the transposase sequence, and a second plasmid encoding Tol2, the promoter, the neomycin resistance gene, a gene encoding for Epac-S^H250^, and a second Tol2 sequence. HeLa iCas9 and Cos7 iCas9 cells were seeded onto 6-well plates at approximately 10% density and transfected the following day. A mixture of 1 μg of each plasmid with 6 μl of FuGENE6 (Promega, #E2692) reagent was added to 200 μl of serum-free DMEM and incubated for 30 minutes. Afterward, the mix was added to the cells and further incubated for 48 hours. Subsequently, the cells were subjected to G418 (200 μg/mL, Sigma-Aldrich, G418-RO) selection.

A monoclonal cell line was generated by preparing a highly diluted cell suspension and seeding it into 96-well plates. Individual colonies were manually inspected, and those derived from single cells were selected for further analysis. Clones were assessed for fluorescence brightness (mTurquoise and PA-mCherry) and evaluated for the largest FLIM span when stimulated with 25 µM forskolin (Sigma Aldrich, #F3917).

### Generation of individual KO cell lines and testing

#### gRNA design and cloning

For generating the Cos7^H250^ ADRB2 polyclonal KO cells, gRNA sequences (20 bp) were selected from the Gattinara Library (Addgene #13698642^43^). For generating all the other KO lines in HeLa^H250^ cells (for hit validation of the screens), guide sequences (20bp) enriched during the primary screens were utilized.

Forward and reverse oligonucleotides containing appropriate overhangs were phosphorylated, annealed, and ligated into the BsmBI-digested pXPR_050 vector (Addgene #96925). Ligated plasmids were transformed, and successful clones were confirmed via Sanger sequencing. Plasmids for transfection were purified (Invitrogen, Thermofisher, #K210004) according to the manufacturer’s instructions. All gRNA and primer sequences are detailed in Supplemental File S1.

#### Lentivirus production and transduction

Lentiviral particles were produced by co-transfecting 40 × 10^6^ Hek-293 (CRL-1573) cells seeded in 10 cm plates with 300 ng/mL gRNA plasmid, 275 ng/mL psPAX2, and 75 ng/mL pMD2.G using 3 µg/mL polyethylenimine (PEI) in Opti-MEM (Gibco, Thermofisher, #31985070). The culture medium was replaced 24 hours post-transfection. Virus-containing supernatants were harvested at 72 hours, filtered through a 0.45 µm syringe filter, and either used immediately for transduction or stored at −80 °C.

To generate individual KO lines, target cells were transduced with high-titer supernatant. Selection was initiated 24 hours post-transduction using 2 µg/mL puromycin (InvivoGen, #ant-pr-5b), and gene editing was induced by the addition of 2 µg/mL doxycycline (Sigma Aldrich, #D9891). Puromycin selection was maintained for 72 hours, while doxycycline treatment was continued for a total of 7 days to ensure robust editing.

#### Making a monoclonal KO cell line

Monoclonal KO cell lines were generated by preparing a highly diluted cell suspension from the polyclonal KO population and seeding it into 96-well plates. Single-cell-derived colonies were identified by manual inspection and expanded for further analysis. Clones were pre-screened for fluorescence intensity by making use of the mTurquoise reporter present.

#### Characterization of KO cell lines

For phenotypic testing, the KO cell lines (both polyclonal and monoclonal) are seeded on coverslips a day before imaging, before subjecting them to their respective validation assays. For genotypic testing, genomic DNA (gDNA) was extracted from both polyclonal and monoclonal populations using the gDNA Isolation Kit (Meridian Biosciences, #BIO-52067). The targeted genomic loci were amplified by PCR using specific primers (Supplemental File S1) and analysed by Sanger sequencing. Editing efficiency and Indel distributions were quantified using the Tracking of Indels by Decomposition (TIDE) software tool^44^. Sequence files are available in Supplementary Sequencing data.

### Performing FRET-FLIM experiments

FLIM experiments were conducted on a Leica Stellaris 8 FALCON confocal microscope, equipped with LAS-X version 4.8.2.29567^45^. The system was equipped with a stage-top incubator maintained at 37°C with 5% CO₂. As our sensor incorporates a tandem dark Venus acceptor, it allows capturing a large part of the donor emission spectrum while minimizing interference from acceptor emission. Detailed imaging settings are outlined in Table 1. The recorded TCSPC photon arrival time histograms and phasor plots showed multi-exponential decay, suggesting a superposition of (at least) two FRET states. Therefore, following acquisition, the photon arrival times were fitted to a double-exponential reconvolution function with fixed lifetimes *τ*_i_ of 0.6 ns and 3.4 ns, representing the previously characterized high-FRET state and low-FRET state, respectively. They were exported from LAS X as *ImageJ TIF* files with 0.001 ns per gray value and used in our analysis macro, FAST-HIPPOS.

**Table 1.** Imaging settings used for signal detection from the Epac^H250^ sensor, PA-mCherry, and PA-JF646.

| Imaging conditions | Epac sensor | PA-mCherry | PA-JF646 |
| --- | --- | --- | --- |
| Lasers used | 440 nm, 448 nm @ 40 MHz | 561 nm | 638 nm |
| Detector(s) | HyDX2 for FLIM [460 nm – 552 nm] | 575 – 660 nm | 650 – 690 nm |
| Bidirectional scanning | Yes | Yes | Yes |
| Zoom | 0.75 | 0.75 | 0.75 |
| Pinhole | 5.00 | 5.00 | 5.00 |
| Scan format | 512 x 512 | 512 x 512 | 512 x 512 |
| Scan speed | 400 | 400 | 400 |
| Objective | 20x 0.75 NA dry | 20x 0.75 NA dry | 20x 0.75 NA dry |
| AFC used | Yes | Yes | Yes |

For experiments involving biochemical stimulation, glass-bottom dishes (Willco Wells, #HBSB-3522) are filled with 2 mL FluoroBrite (FB) medium (Gibco, Thermofisher, #A1896702) prior to imaging and are transferred to the microscope stage. First, ∼1-2 minutes of basal cAMP levels are measured, followed by stimulation with 1 µM or 40 nM isoproterenol (Sigma Aldrich, #PHR2722), a β-adrenergic receptor agonist, and then calibration with 25 µM forskolin. Detailed settings are also available as metadata saved in the Leica .lif files.

### Photoactivation of the H-NS-PA-mCherry

Prior to imaging, culture medium was replaced with FB medium in glass-bottom dishes containing Cos7^H250^ cells. Photoactivation was performed using an optimized protocol. Single cells were activated with a 405 nm diode laser set at 0.215 mW laser power, using a 20X 0.75 NA dry objective and 48x zoom. Two rounds of excitation were applied with the Leica resonant scanner at 8000 Hz, bi-directional imaging, yielding a pixel dwell time of 0.369 µs, in a 64×64 image format with 64x accumulate. A slight defocus of +6 µm was also applied to prevent excessive bleaching. For bulk-cell maximal photoactivation, cells were seeded in 6-well plates, and a benchtop 395 nm LED (230 mW/cm^2^) that fitted exactly on top of each well was used for 6 minutes. Settings used for readout of PA-mCherry are outlined in Table 1.

### Loading and photoactivation of PA-JF646

Prior to imaging, cells were incubated with 5 mM NH_4_Cl for 1 min, then replaced with FB medium containing 1 µM PA-JF646 dye (Tocris Bioscience, #6150). Cells were incubated for 30 min at 37°C and then washed once with PBS and once with FB medium. Dishes were subsequently filled with 2 mL FB medium and transferred to the microscope stage for imaging.

Photoactivation of PA-JF646 in single cells was performed with an external 355 nm pulsed laser (Spectra-Physics Explorer One 355-1) set at 50 kHz repetition rate, for 1.66 seconds per position with an average power of 1.7 µW. The laser beam was collimated by a 2-lens telescope system to a beam waist of ∼5 mm at the back of the objective, thereby underfilling the backfocal plane. Additionally, the beam was slightly defocused, in such a way that the diameter of the spot in the sample plane was about 8 µm, slightly smaller than the size of a cell nucleus, enabling single-cell activation without affecting neighbouring cells.

The position of the focal plane was actively stabilized using the Leica Auto Focus Control (AFC) to prevent focal drift during photoactivation of multiple regions. Bulk cells were maximally photoactivated by seeding them in 6-well plates and using a benchtop 395 nm LED (230 mW/cm^2^) _f_or 6 minutes that exactly fit on top of each well. Settings used for readout of PA-JF646 are outlined in Table 1.

### Performing the simulation experiment

For the simulation experiment, Cos7^H250^ ADRB2 polyclonal KO cells were pre-labeled with 1 µM PA-JF646 dye, photoactivated, and then trypsinized. Approximately 7% of these KO cells were then seeded in a dish containing unlabeled wild-type (WT) cells (75% confluency). The mixed population was allowed to settle for 6–7 hours prior to imaging. Time-lapse FLIM imaging was performed for 30 min, with stimulation using 40 nM isoproterenol followed by calibration with 25 µM forskolin.

Given that we mixed ∼7% polyclonal KO cells with a 41% phenotypic penetrance, we expected a ∼2.9% hit rate, supplemented by a 1.8% false-positive rate from the WT population (totaling 4.6%). The slight surplus actually observed (0.6 %) likely results from minor non-homogeneous cell distribution on the imaging dish during seeding.

### Preparing photoactivated cells for FAC sorting

Once imaging and photoactivation of all samples were complete, the samples were processed together for FACS. Samples with the PA-JF646 dye were prepared and handled in the dark using aluminum foil to avoid even minimal background activation. After trypsinization and treatment with trypsin inhibitor, cells were pelleted by centrifugation at 1400 rpm for 4 min, resuspended in FB medium, and filtered through a cell-strainer (Corning, #352235) to obtain a single-cell suspension. Samples were kept on ice until sorting. Cell sorting was performed on a BD FACS Fusion cell sorter (BD Biosciences). Forward- and side-scatter parameters were used to exclude debris, doublets, and dead cells. Fluorescent signals were acquired using the following laser/filter combinations: PA-mCherry was excited with a 532 nm laser and detected with a 610/20 nm filter; mVenus was excited at 488 nm and detected at 530/30 nm; and PA-JF646 dye was excited with a 640 nm laser and detected at 670/30 nm.

Positive and negative controls were included for gating: positive controls consisted of cells maximally photoactivated with a benchtop 395 nm LED (for both PA-mCherry and PA-JF646). Negative controls for the dye were incubated with 1 µM dye but not photoactivated, whereas negative controls for PA-mCherry were non-activated cells.

### Dolcetto library benchmark experiment

Cos7^H250^ WT cells were transduced separately with Dolcetto library set A (57,050 gRNAs) and set B (57,011 gRNAs). A total of 10^6^cells were used for transduction, with 128 µL of set A (MOI = 0.3) and 256 µL of set B (MOI = 0.4). After transduction, cells were subjected to puromycin selection (2 µg/mL) for three days to ensure transduction efficiency. Post-selection, set A cells were seeded in a 6-well plate, while set B cells were seeded in a 10 cm dish. Two wells of set A cells were photoactivated using a benchtop 395 nm LED and subsequently mixed with the non-photoactivated set B cells in the 10 cm dish (∼1:4). The resulting mixed population was prepared for FACS (as described in section *‘Preparing photoactivated cells for FAC sorting’*), and samples of 200, 500, 2,000, 5,000, and 10,000 activated red cells were sorted directly into PCR tubes containing 20 µL of lysis mix (prepared by mixing 200 µL lysis buffer (Viagen, #301-C), 200 µL ultra-pure nuclease free water (ThermoFisher, #2844363), and 2 µL Proteinase K (Viagen, #501-PK) and were centrifuged at 1,100 rpm for 2 min to pull down remnants, if any.

Lysis was performed using a thermal cycler: 3 h 15 min at 55 °C followed by 45 min at 80 °C to inactivate Proteinase K. Lysates were used directly in the first round of PCR (PCR1), with four parallel reactions per sample, each using 5 µL of lysate. PCR1 conditions are detailed in Table 2. A unique forward primer was used for each sample (Supplemental File S2). This primer introduces an Illumina Read 1 sequencing adapter and a sample-specific barcode. The barcode allows us to index each sample separately and deconvolve the sequence reads after the pooled sequencing run. PCR1 products for each sample were pooled, and two replicate PCR2 reactions (5 µL input each) were performed to add flow cell adapters required for Illumina sequencing. PCR2 conditions are listed in Table 2. Together, the two rounds of PCR amplified the low DNA input to sufficient amounts for sequencing. To account for sample-to-sample variation in DNA yield, normalization was performed. A 10 µL aliquot of each PCR2 product was resolved on a 1% agarose gel to verify amplicon size and assess DNA quantity (Fig. S1.4 (i), (ii)). Band intensities were quantified to determine relative DNA concentrations and used to pool samples in defined ratios. The pooled sample was resolved again on a 1% agarose gel, and the desired product was excised and gel-purified with a purification kit (Meridian Biosciences, #BIO-52060). The final gel-purified product was submitted for next-generation sequencing on the Illumina MiSeq platform.

**Table 2.** PCR1 and PCR2 conditions.

|  |  |  |  |  |  |  |  |  |
| --- | --- | --- | --- | --- | --- | --- | --- | --- |
| PCR 1:<br>(Product size<br>~280 bp) | 5 μL – Lysate<br>1 μL (10 μM) – FP<br>1 μL (10 μM) – RP<br>2 μL – MyTaqRedMix<br>18 μL – milliQ | 98°C –<br>30 s | 98°C –<br>30 s | 60°C –<br>30 s | 72°C –<br>1 min | 72°C –<br>5 min | Hold<br>at 4°C | 35<br>cycles |
| PCR 2:<br>(Product size<br>~350 bp) | 5 μL PCR1 pooled<br>product<br>1 μL (10 μM) – FP<br>1 μL (10 μM) – RP<br>25 μL – MyTaq2Red<br>18 μL – milliQ | 98°C –<br>30 s | 98°C –<br>30 s | 60°C –<br>30 s | 72°C –<br>1 min | 72°C –<br>5 min | Hold<br>at 4°C | 16<br>cycles |

Note that we found it to be crucial to use Taq polymerase (MyTaqRedMix, Meridian Biosciences, #BIO-25044) because of its robust performance with crude, low-input samples. However, its high sensitivity to contamination necessitated strictly sterile sample handling, tips, pipettes, and reagents. Water controls and all reaction steps were included in every run to monitor for contamination. Alternative polymerases, such as Phusion (New England Biolabs, #M0530L), failed to amplify DNA under these conditions.

### MTT (Toxicity) assay

To investigate cytotoxicity, 5×10^4^, 2×10^4^, or 1×104 Cos7 WT cells were seeded in 96 wells and incubated for 4 hours, 24 hours, or 48 hours, respectively, with 1 nM, 10 nM, 100 nM, or 1 µM of the PA-JF646 dye. A control with DMSO incubation was taken along, as well as a control with medium and dye without cells. After the dye incubation, MTT labeling reagent (end concentration of 0.5 mg/ml) was added to each well and incubated for another 2 hours. Then, the medium was removed, and 100 µL DMSO was added. After all formazan crystals were dissolved, absorbance was read at 540 nm. MTT reduction is measured as the absorbance at 540 nm, normalized to the DMSO control.

### Performing the screens

#### Pilot screen and dynamic OPS - Lentiviral library designing, cloning, production, titer calculation, sequencing, and verification

##### Design, cloning, and production

Guide sequences were designed using the Broad GPP portal, and oligos were designed with 15 bp overhangs to facilitate cloning. The oligo pools were ordered from Twist Biosciences and cloned into the pXPR_050 vector using Gibson assembly as described in this paper^46^. To generate lentivirus containing the gRNA library, the same protocol as generating lentivirus with individual gRNAs was used (See section *‘Lentivirus production and transduction’*).

#### Titer calculation

To determine the titer, 150,000 Hela^H250^ cells were seeded in a 6-well plate and transduced with 5 different volumes of the virus (512 µL, 256 µL, 128 µL, 64 µL, 32 µL, 16 µL). An untransduced control was also taken along. After 24h of transduction, the medium was replaced with fresh growth medium. At 48h after transduction, the cells are put under puromycin (2 µg/ml) selection for 2 days. Surviving cell fractions were used to calculate the MOI based on the fraction of resistant colonies. The MOI was determined using the equation:

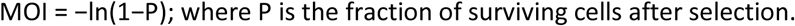

##### Sequencing and verification

10 ng plasmid DNA was used as a template to amplify the gRNA sequences. The PCR strategy was similar to that described in the section *‘Dolcetto library benchmark experiment’*; the only difference was that 25 cycles were used for PCR1. The PCR products were sequenced on the Illumina MiSeq platform. The sequence reads were mapped to the custom gRNA library, and the gRNA recovery and distribution were computed in RStudio. All 1272 gRNA sequences were detected, and the distribution of the gRNAs (79.2% within 2-fold difference from the median and 99.9% within 10-fold difference from the median) was similar to that observed for other libraries (Supplemental File S3).

#### Pilot screen and dynamic OPS – Preparing the cells

For both screens, cells were prepared similarly. The screens were conducted in three biological replicates. For the dynamic OPS, each replicate was performed on different days using the same cell line and identical protocols. For the pilot screen, all replicates were performed on the same day. Steps are outlined here for *one* replicate.

On day 1, HeLa^H250^ cells were seeded into five (seven for the pilot screen) 10 cm dishes, each containing 500,000 cells. Cells were seeded as follows:

Plate 1 = Sample REP1 (Transduced + Puromycin)
Plate 2 = Control (Transduced – Puromycin)
Plate 3 = Control (Untransduced + Puromycin)
Plate 4 = Imaging Control (Untransduced – Puromycin)
Plate 5 = For counting cells the next day
Plate 6 and 7 = Sample REP2, Sample REP3 (Transduced + Puromycin) [only for pilot screen]

On day 2, the plate designated for cell counting was used to estimate cell numbers, and the other plates were transduced with 128 µL of lentivirus (MOI = 0.3) carrying the library. On day 3, cells were passaged at 20% confluency, reseeded into new plates, and subjected to 2 µg/mL of puromycin selection while editing was induced by adding 2 µg/mL of doxycycline. Doxycycline was also added to the Imaging control to keep the same culture conditions. On day 6, puromycin selection was discontinued, but doxycycline was maintained. Only the imaging control was split at 30% confluency, while transduced plates were allowed to proliferate.

The day before the screen (Day 8), 10 glass-bottom dishes were seeded with 150,000 cells each from the transduced sample, and 2 dishes were seeded with 150,000 cells each from the imaging control. Additionally, 2 dishes were seeded with 10^6^ cells each from the transduced sample to serve as the reference (t_0_) controls. Penicillin-streptomycin (ThermoFisher #15140122) was added to all samples and controls to prevent contamination. On the day of the screen (Day 9), cells were visually inspected for health and confluence before proceeding.

#### Pilot screen and dynamic OPS - Imaging the cells

Prior to imaging, the cells are loaded and incubated with PA-JF646 as described earlier (See section ‘*Loading and photoactivation of PA-JF646’*). Following that, they are imaged as described earlier (See section ‘*Performing FRET-FLIM experiments*’). The details of stimulation, calibration, number of tiles, and number of cells imaged are summarized in Table 3.

**Table 3.** Summary of dynamic OPS and pilot screen.

| Screen type | Dynamic OPS | Pilot screen |
| --- | --- | --- |
| Phenotype screened for | Low cAMP responders, Transient cAMP responders | Cells with high fluorescence lifetime levels |
| Library type | Custom-made; 318 genes, 1272 guides | Custom-made; 318 genes, 1272 guides |
| MOI | 0.3 | 0.3 |
| Number of tiles imaged | 240 | 304 |
| Total number of frames imaged | 10 | 2 |
| Imaging time | 1 hour | 12 minutes |
| No. of glass-bottom dishes screened | REP 1; REP 2; REP 3 → 5; 4; 4 | REP 1; REP 2; REP 3 → 4; 4; 4 |
| Total time taken (3 REPS) | 3 days | 1 day |
| No. of cells screened | REP 1; REP 2; REP 3 → 129,136; 123,710; 115,368 | REP 1; REP 2; REP 3 → 177,602; 160,842; 169,025 |
| Coverage | REP 1; REP 2; REP 3 → 101x; 97x; 91x | REP 1; REP 2; REP 3 → 136x; 123x; 130x |
| No. of hits (cells) activated | REP 1; REP 2; REP 3 → 4356; 4898; 4593 | REP 1; REP 2; REP 3 → 913; 1041; 889 |
| No. of cells sorted | REP 1; REP 2; REP 3 → 3375; 1735; 2017 | REP 1; REP 2; REP 3 → 292; 510; 379 |
| Stimulation type | 1 μM isoproterenol (at end of Frame 2), 25 μM forskolin calibration (at end of Frame 8) | None |
| Criteria for hit selection | Response lifetime difference with the cell's own baseline in a 3-frame window should be lower than 0.4 ns + Baseline to calibration lifetime difference is more than 0.30 ns + Absolute baseline to average baseline difference is less than 0.30ns | Any cell with a baseline lifetime value > 2.6 ns |
| Photoactivation method | PA-JF646 and external laser 355 nm | PA-JF646 and external laser 355 nm |
| Time taken for FAST-HIPPOS analysis | ~8 minutes | ~5 minutes |

#### Pilot screen and dynamic OPS - Image analysis and hit detection

Tiles are exported as individual tiles rather than in a merged format. They are then stitched together using a Fiji script that in turn incorporates the Grid/Collection Stitching plugin^47^ (Stitch_tiles macro; https://imagej.net/plugins/fast-hippos). This is deliberately done without the calculation of overlap in order to preserve stage positions. XY-Stage coordinates of image tiles are extracted from the OME metadata of the .lif file and stored in the metadata of the stitched .tif file, together with the grid layout, the tile size, and the tile overlap percentage. The stitched image is then fed into the FAST-HIPPOS macro. Here, an output folder is defined, and cell segmentation and hit-detection criteria are added. The analysis time and hit-detection criteria for both screens are outlined in Table 3 below. Among many output graphs and figures, the macro also generates a file with the stage coordinates of the hit cells (.rgn files in XML layout) (Supplemental Material 2). This is then imported into LASX Navigator to start the photoactivation regime.

#### Pilot screen and dynamic OPS - Photoactivation of hit cells

Photoactivation of identified hit cells was performed as described earlier (See section ‘*Loading and photoactivation of PA-JF646*’).

Following photoactivation, glass-bottom dishes were stored in an incubator to maintain cell viability. Once imaging and photoactivation of all samples were complete, the samples were processed together for FACS. Samples containing the PA-JF646 dye were prepared and handled in the dark, using aluminum foil to prevent background activation.

#### Pilot screen and dynamic OPS – Preparing and FAC sorting hit cells and sample preparation for NGS

Photoactivated samples and FACS controls were prepared and sorted as described earlier (See section ‘*Preparing photoactivated cells for FACS sorting*’). Sorting, cell lysis, and sample preparation for NGS were also performed as described earlier (See section ‘*Dolcetto library benchmark experiment*’).

#### Pilot screen and dynamic OPS - Bioinformatics analysis

With the NGS data, both CRISPR screens (pilot and dynamic OPS) were analysed using the following approach: as a first step, we normalized sequence counts to account for variations in sequencing depth between samples. We implemented a normalization method analogous to that used by DESeq2^48^, but with an important adaptation. Specifically, we calculated a relative total size factor by taking the mean over all sample totals and dividing each sample total by this mean. Each count value within a sample was then divided by the corresponding size factor for that sample, resulting in equalized totals across all samples. We chose a size factor based on total counts rather than the median counts, which DESeq2 uses, because some samples exhibited massive dropout in the sorted population, which would have resulted in a median value of zero for those samples, making normalization impossible.

The statistical analysis was performed in two sequential steps. First, we conducted a differential analysis between the sorted and reference populations using DESeq2, with the normalized counts as input, skipping the DESeq2 normalization step. Second, we performed an enrichment analysis on the gRNAs using MAGeCK’s Robust Rank Algorithm (RRA)^49^ using the results from the DESeq2 analysis as input. Because the RRA analysis in this case is about positive selection, we sorted the DESeq2 results on the DESeq2 test statistic from high to low. For the RRA analysis, we set the threshold for ‘good gRNAs’ (controlled by the maxPercentile parameter) based on our DESeq2 results. Specifically, we calculated this threshold as the percentage of gRNAs that showed both a positive test statistic (corresponding to a positive log2 fold change) and a p-value ≤ 0.1 relative to the total number of gRNAs in the dataset.

For hit selection, we used an FDR calculated by RRA ≤ 0.1 and a median log2 fold change ≥ 1 across all gRNAs targeting a gene.

#### Pilot screen and dynamic OPS - Hit validation

After identification of the hit genes in both screens, validation experiments were performed. For that, the following KO cell lines were made using the protocol described earlier (See section ‘*Generation of individual KO cell lines and testing*’):

HeLa^H250^RAPGEF3/Epac1 polyclonal KO cell line
HeLa^H250^ ADRB2 KO, HeLa^H250^ GNAS KO and HeLa^H250^ ADCY6 KO (all monoclonal)

For phenotypic validation of all KO cell lines, cells were subjected to the same imaging assay used during their respective screens. The data was analysed using FAST-HIPPOS.

#### ELISA Assay

cAMP ELISA assay was performed using a Cyclic AMP XP^®^ Assay Kit (#4339, Cell Signaling Technology) according to the manufacturer’s protocol. Cells were seeded in a 48-well plate and grown overnight. All experiments were performed in biological triplicate. No stimulations were necessary as we were measuring basal levels of cAMP, except for the forskolin control. The following samples were tested: HeLa^H250^ WT (Control)

Hela^H250^ WT treated with 25 µM forskolin
HeLa^H250^ RAPGEF3/Epac1 polyclonal KO

### Statistical Analysis

A custom R script written in RStudio (version 2023.03.0) was used for data visualization (codes available at Zenodo repository - https://doi.org/10.5281/zenodo.20068508). Briefly, FAST-HIPPOS generates lifetime data of every cell imaged per timepoint, which are exported as .csv or .tsv files. The R script runs through the .csv or .tsv file, to determine baseline lifetime values for each cell from the first 10 time points. The script allows setting criteria for the inclusion of each individual cell based on outlier rejection and the variance of time points within the baseline. Typically, between 1 and 15% of cells were rejected, often because of poor segmentation and/or detached cells that affect the validity of the data.

For lifetime traces, since a typical FOV contains up to 300 individual cells, we choose to plot representative individual traces, picked randomly from all cells that pass the inclusion criteria. To still fully capture the range and variability of the data, we also included a summary dotplot of the relevant data (marked with grey dotted lines) from all included cells, displayed next to the traces and at the same Y axis scale. Dotplots also show boxplots with median value (horizontal black line), middle 50% of values (boxes), and 1.5 times the interquartile range (whiskers). For histograms, all the cells are included in the plots.

Testing for statistical significance of relevant differences was carried out by unpaired one-tailed t-tests, and all were performed using GraphPad Prism (version 10.3). In the main text and legends, significances are marked as: *\*p < 0.05; **p < 0.005; ***p < 0.0001;* ‘ns.’ = not significant.

### Plasmids used

The gRNA cloning vector pXPR_050 (#96925) and the Edit-R Inducible Lentiviral Cas9 vector (#NC1606271) was purchased from Dharmacon.

### Data and software availability

All raw data sets, analysed data and R scripts are available on Zenodo repository – https://doi.org/10.5281/zenodo.20068508.

FAST-HIPPOS is available at: https://imagej.net/plugins/fast-hippos

## ACKNOWLEDGMENTS

We are indebted to Dr. L. Bombardelli (Leiden University) and Dr. O. Kukk (Solis BioDyne) for insightful discussions and hands-on help during the initial phase of the project. The H-NS plasmid was kindly provided to us by Dr. Remus T. Dame (Leiden University) before publication of their manuscript. We are grateful to Daniil Anastasopoulos (NKI, Amsterdam) for helping make the initial design of the H250 construct and to Elke Malzer (NKI, Amsterdam) for all the support with downstream PCR processing. We are also grateful to the NKI FACS and Genomics facility for their technical support with all the experiments.

This study was supported by funding from NWO (TTW 14691 to K. Jalink). Research at the Netherlands Cancer Institute is supported by institutional grants of the Dutch Cancer Society and the Dutch Ministry of Health, Welfare, and Sport. Part of the work has also been enabled by the European Union’s Horizon Europe research innovation program under Marie Skłodowska-Curie Grant Agreement No. 101073507 (flIMAGIN3D – HORIZON-MSCA-DN-2021) and the NL-Bioimaging infrastructure sponsored by NWO National Roadmap for Large-Scale Research Infrastructure (NWO 184.036.012). Additional support was also received from ScreeninC Infrastructure (KWF 12539).

## AUTHOR CONTRIBUTIONS

Conception of the study: K.J. and R.L.B.; design and execution of FLIM experiments: S.M. and K.J.; Designing of the library: S.M.; Generation of the library: R.L.B.; Sequencing of library and Dolcetto experiment: H.J.K. and S.M.; Optimization of H-NS-PA-mCherry: S.M. and K.J.; Optimization of the PA-JaneliaFluor646 – M.v.T., S.M., B.vd.B. and K.J.; Screens: S.M., D.S. and D.K.; Validation of hits from screens: S.M., D.K., and G.Z.; Bioinformatics analysis of NGS data: C.L.; preparation of molecular constructs, construction of stable cell lines, and cell culturing: S.M., J.K., G.Z., D.K. and M.v.T; ImageJ scripts for automatic cell segmentation, extraction of single-cell FLIM traces and hit identification: B.vd.B., K.J. and S.M.; data analysis, statistics, and preparation of figures: S.M., K.J. and B.vd.B.; R scripts for data visualization: S.M. and K.J.; ELISA assays: S.M.; preparation of the manuscript: S.M. and K.J. All authors provided critical input during finalizing of the manuscript.

