## Supplemental Material for "Beyond Static Screens: optical pooled screening of signaling dynamics using time-lapse FLIM"

### SUPPLEMENTAL MATERIAL 1: ADDITIONAL FIGURES

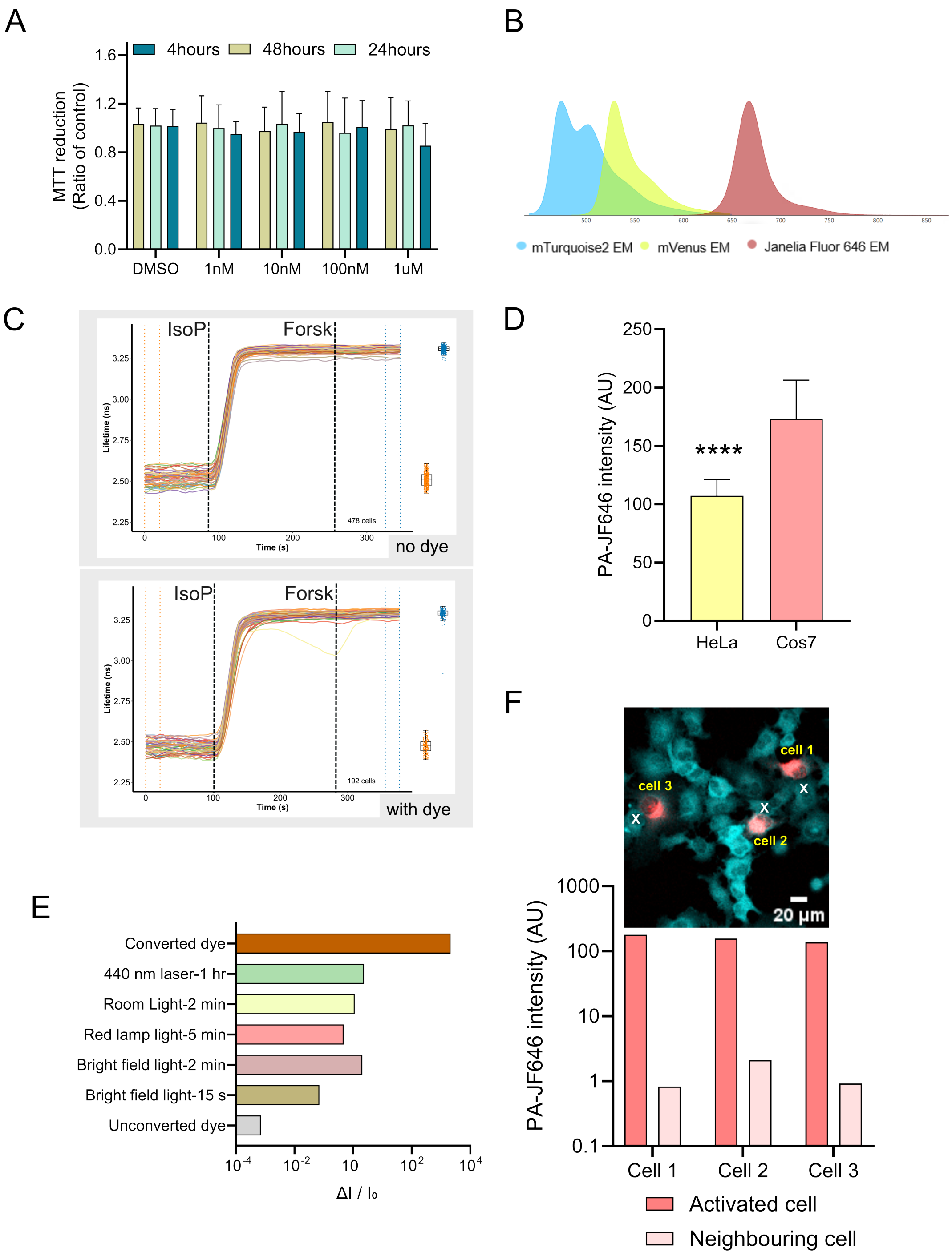


***Figure S1.1. Additional optimization experiments of PA-JF646. (A)*** *MTT assay for PA-JF646 dye cytotoxicity. Cells were incubated with 1 nM, 10 nM, 100 nM, or 1 µM PA-JF646 for 4, 24 or 48 h; no significant differences in viability were found. Statistical analysis was performed using a two-way ANOVA with a multiple-comparison test; $p > 0.05$.* ***(B)*** *Spectral separation of PA-JF646 from Epac-$S^{H250}$ sensor emissions. The PA-JF646 emission peak (~665 nm) is well*

separated from both donor and acceptor peaks. ***(C)*** *Effect of PA-JF646 on cAMP sensor readout. FLIM traces of Cos7$^{H250}$ cells without (top, n = 478) or with (bottom, n = 192) PA-JF646 dye incubation. Orange box plots show baseline lifetimes of all cells in the first 10 frames (orange dotted lines); blue box plots show post-forskolin saturation levels of all cells in the last 10 frames (blue dotted lines). IsoP denotes the addition of 40 nM isoproterenol, and Forsk denotes the addition of 25 µM forskolin.* ***(D)*** *PA-JF646 fluorescence (AU) in Cos7$^{H250}$ cells and HeLa$^{H250}$ cells upon photoactivation with the 355 nm laser (n=33). Intensity differences ($I_{post\ activation} - I_{pre\ activation}$) are shown as mean ± SD of the ROI-averaged pixel values. Statistical analysis was performed using an unpaired, one-tailed t-test. ****p < 0.0001.* ***(E)*** *Spurious photoactivation of PA-JF646 under different light conditions. Change in fluorescence intensity reported as ($\Delta I/I_o$); x-axis is plotted in log(10) scale. Fully converted dye shows maximal signal. Measurements are based on a single experiment. Shown are mean intensities over the entire field of view.* ***(F)*** *Neighbouring cell activation during PA-JF646 activation. The image shows 3 target cells, and the closest adjacent cell is marked with X; the bar graph shows their respective intensity differences, $I_{post\ activation} - I_{pre\ activation}$.*

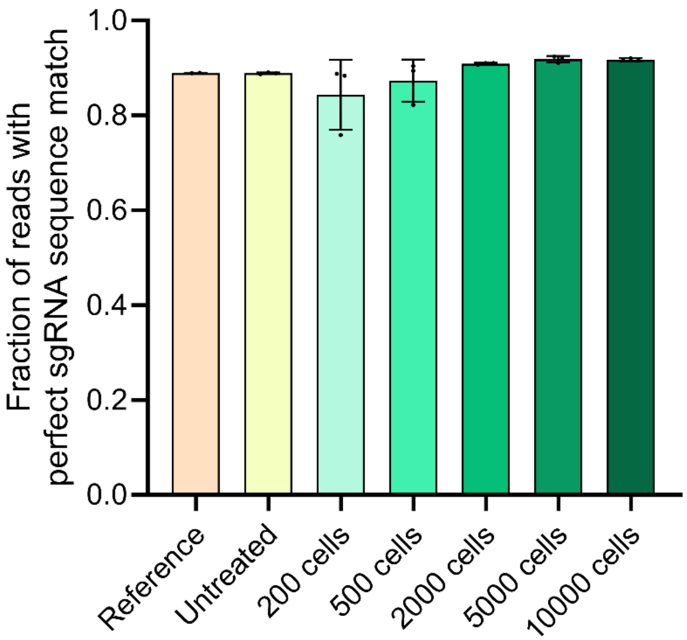


***Figure S1.2. Assessment of gRNA sequence fidelity after sample preparation using Taq polymerase.*** *Fraction of sequencing reads that perfectly match reference gRNA sequences in the library. "Reference" and "Untreated" are samples (the whole setA library) prepared in an independent experiment using high-fidelity polymerases. Samples labeled 200 to 10,000 cells were processed and prepared using our benchmarked method as described in M&M for NGS. Bars represent mean ± SD from triplicate preparations. Comparable fractions of perfect matches across all conditions indicate that our method does not introduce additional mutations during amplification.*

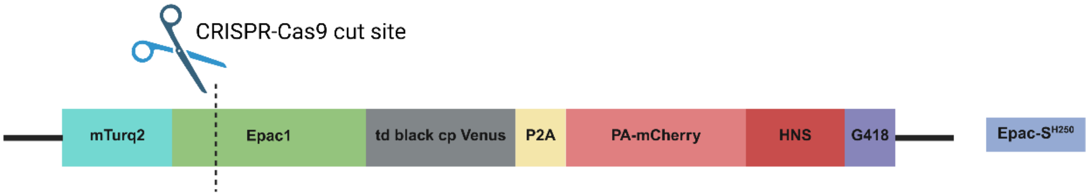


***Figure S1.3.*** *RAPGEF3/Epac1 cut site that is expected to take out most of the Epac1, as well as the FRET acceptor in the reporter construct Epac-S*$^{H250}$.

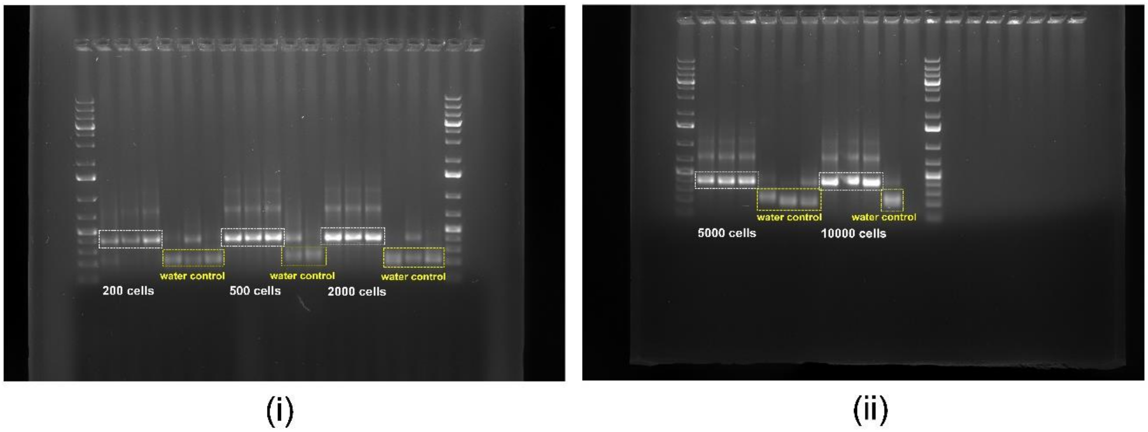


***Figure S1.4. (i)*** *1% Agarose gel ran with PCR 2 samples containing samples with 200, 500 and 2000 cells (in triplicates) and their respective water controls* ***(ii)*** *1% Agarose gel ran with PCR 2 samples containing samples with 5000 and 10000 cells (in triplicates) and their respective water controls. In both gels, left- and rightmost lanes are 1 kb ladder.*

**SUPPLEMENTAL MATERIAL 2: DIFFERENT FUNCTIONALITIES OF FAST-HIPPOS**

FAST-HIPPOS: FLIM Analysis of Single-cell Traces for Hit Identification of Phenotypes in Pooled Optical Screening is a collection of macros in Fiji[1] to analyse time-lapse FLIM data and to automatically identify hit cells and subsequently drive their photoactivation (Fig. S2.1). The script can load time-lapse FLIM data directly from Leica .lif files, Lambert Instruments .fli files, and exported (ome) .tiff files. Simple intensity data and ratio imaging data are also supported. Optional preprocessing steps include drift correction and bidirectional scanning phase mismatch correction.

Using these data, cell segmentation can be performed with CellPose[2] in Fiji (https://github.com/BIOP/ijl-utilities-wrappers) on the summed-intensity projection of a subset or all of the time-lapse images (Fig. S2.1A). Based on the obtained segmentations, temporal lifetime traces are extracted for individual cells by intensity-weighted averaging the pixels belonging to each cell (Fig. S2.1B). Lifetime trace data can be represented in a variety of ways: as time traces, graphs, tables, density graphs, kymographs, and more. Optional outputs include (smoothed) lifetime-intensity overlay movies (Fig. S2.1A and Supplemental movie 1 and 2), (time-lapse) lifetime histograms, and (time-lapse) scatter plots of cell lifetimes against various parameters (Fig. S2.1C and Supplemental movie 3 and 4).

If applicable, the script can automatically determine the time points of stimulation and calibration based on steep inclines in the average trace, and can use this information to subdivide traces into 'baseline', 'response', and 'calibration' segments. The data from a user-selectable range of time frames can then be used to call hit cells through a multi-parameter scan of the data traces, with user-defined selection criteria, e.g., baseline lifetime, maximum response lifetime, average lifetime in a defined time range after stimulation or before calibration, etc. (Fig. S2.1D). Alternatively, hit cells can be selected graphically by simply drawing a ROI in time traces, a sorted kymograph, or the overlay movie (see Supplemental movie 5). The detected hits are displayed in a time traces graph, outlined on the lifetime-intensity overlay movie for inspection (Fig. S2.1E), and are shown in density graphs, kymographs, and (dynamic) time-lapse histograms (Fig. S2.1F). In 'inspection mode', data from different graphs are visually coupled, i.e., clicking on a particular trace will highlight the corresponding cell and vice versa (Supplemental movie 6), enabling quick examination of the data.

The (X, Y) stage coordinates of hit cells are saved as Leica .rgn files, which can be directly imported into LAS X Navigator (Fig. S2.1G) for photoactivation. For our screens, we imaged large regions (e.g., 200 tiles ~30,000 cells) without overlap. Image stitching is performed in FIJI to ensure the correct pixel size to prevent image distortion and preserve stage coordinates.

To accelerate analysis, FAST-HIPPOS makes extensive use of GPU acceleration[3]. A detailed description of the analysis pipeline and the script can be found at https://imagej.net/plugins/fast-hippos.

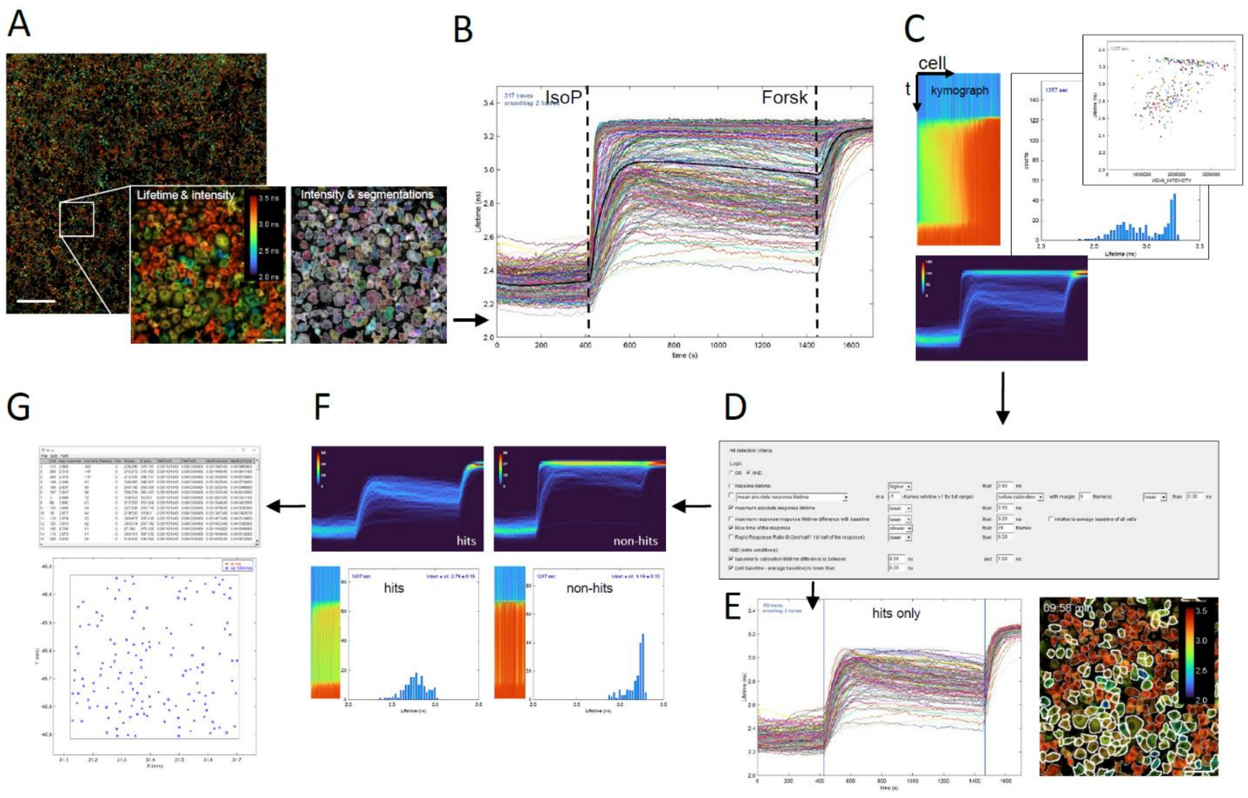


***Figure. S2.1. Time-lapse FLIM analysis and hit detection Fiji macro.*** ***(A)*** *Stitched multi-tile intensity image overlay with the fluorescence lifetime image. The lifetime image can be either average photon arrival times ('Fast FLIM') or calculated average lifetimes from two lifetime components. Ratiometric images can also be used. Scale bar: 1000 µm.* ***Inset:*** *Left – Zoom-in of lifetime image of cells after stimulation with isoproterenol; Right – Zoom-in showing cell segmentation as outlines. Scale bar: 100 µm.* ***(B)*** *Lifetime time traces of all 298 segmented cells in a single tile of Cos7 cells (a 1:1 mix of ADRB2 KO and WT cells). Cells are stimulated with 40 nM isoproterenol (IsoP), followed by a calibration with forskolin (Forsk). Stimulation and calibration time points may be automatically calculated from the average of all traces (thick black line) or entered manually.* ***(C)*** *The script generates several informative visualizations. Shown are a 'kymograph' representation of fluorescence lifetime vs time in all cells, sorted on response magnitude (top-left); a normalized data density graph (2D histogram of lifetime vs time, with cell counts in false color), convenient when analysing thousands of cells (bottom); and (optionally animated) histograms of the fluorescence lifetimes over time (middle) and scatterplots of lifetime vs secondary parameters like cell intensity, cell area or intensity of an additional fluorescence channel of choice (top-right).* ***(D)*** *Hit selection dialog showing various optional criteria. These can be used separately or combined, allowing to screen for a large variety of dynamic phenotypes.* ***(E)*** *Based on the criteria, hit cells are determined and visualized through a 'hits only' lifetime traces plot (left). Blue vertical lines indicate the chosen hit detection time window. The hit cells are outlined with white lines on the lifetime-intensity overlay image (right).* ***(F)*** *Additional visualizations of hit cells (left) and non-hit cells (right), in density graphs, response-sorted kymographs, and time-lapse histograms.* ***(G)*** *Microscope stage coordinates of the hit cells are shown in tables and a 2D graph. The script also generates a .rgn file with positions, which can be directly read by LAS X Navigator, for e.g. photoactivation, high-resolution imaging, etc. For visualization of animated plots, see the website (https://imagej.net/plugins/fast-hippos).*

### SUPPLEMENTAL MATERIAL 3: OPTIMIZING HIGH-THROUGHPUT FLIM RECORDING

**FLIM detection.** In time-lapse screening applications, large numbers of cells must be imaged in a photon-efficient manner to avoid photodamage. Time-correlated single photon counting (TCSPC) is the most photon-efficient manner to detect lifetimes, but the photon count rate of conventional TCSPC instruments must remain below one photon per 10-20 laser pulses *in the brightest cell* in order to avoid serious lifetime errors due to so-called photon pile-up effects[4]. In this study, we use the Leica Stellaris Falcon system, which utilizes a detection scheme with much higher count rates, able to collect usable data at up to 3 photons per pulse. To avoid lifetime bias, laser power was adjusted to have a maximum of 2 photons per pulse, using 456-532 nm emission. Optionally, two FLIM detectors (HyD2 and HyD4) can be combined and tuned side-by-side to collect the mTurquoise2 emission together. In that case, the bandwidth of HyD4 (495-532 nm) was adjusted to give the same count rate as HyD2 (456-490 nm), and the two detectors were assigned to the same lifetime channel in LAS-X (FLIM expert settings) so as to double the count rate. For consistency with earlier experiments, that configuration was not used in the two screens.

**Screening speed.** To screen large numbers of cells in time-lapse mode, a trade-off must be made between the spatial resolution of the cell images and imaging speed. A 2.5x objective will cover large numbers of cells, and thus, a lower number of imaging tiles is necessary to cover the same area as a 40x objective. However, higher magnification objectives typically have higher numeric aperture (N.A.), with higher Photon collecting Efficiencies (P.E. scales with N.A. squared) and higher optical resolution (resolution scales linearly with N.A.). However, for this FLIM screen, the spatial resolution arguably is not a primary concern because we collect data from a cytosolic, freely diffusing FRET sensor.

We therefore tested 4 objectives for screening: a 40x 1.1 NA water immersion objective, a 20x 0.75 NA dry objective, a 10x 0.4 NA objective, and a 2.5x 0.07 NA objective. We documented the speed to cover an area corresponding to ~80,000 cells (10 x 10 mm), the laser power necessary to obtain 1 photon/pulse on average, using a fluorescent plastic preparation (Chroma). In addition, lenses were tested on monolayers of Cos7$^{H250}$ cells to compare bleach rate, and we minimized the scan format so that cells could still be reliably segmented using Cellpose, and the signal-to-noise ratio in single-cell FLIM quantitation was better than 50. Results are summarized in Table S3.1. Based on this analysis, we chose the 20x objective, zoom 0.75, and a 512x512 image format as the best trade-off. We also noted that the bleach rate was inversely correlated with lens magnification, likely because the point spread function, in particular in the direction of the Z-axis, of small-NA lenses is considerably larger. As a result, the excitation power is spread over a larger focal spot, causing less bleaching due to the non-linearity of the relation between bleaching and laser power.

After optimization, final settings were selected: 20x 0.75 NA dry objective, 0.75x zoom, FOV 760 by 760 µm; 512 lines with pixel dwell time 1.2 µs; bi-directional scanning enabled. The laser was adjusted to give up to 2 photons per count in the brightest cells in each of the detectors, using Leica's fast counting mode. With these settings, single cells occupied on average 250 pixels, and lifetimes detected over the entire cell had a signal-to-noise ratio of, typically, >>50 (two detectors) or >>35 (a single detector).

With these settings, we could image up to 50,000 cells about once every 3.5 minutes, depending on confluency.

| Lens mag. | Format (px) | line speed (Hz) | Tiles | Pix size (μm) | dwell time (μs) | ~photons/cell | duration (s) |
|---|---|---|---|---|---|---|---|
| 10x | $1024^2$ | 300 | 49 | 1.5 | 1.2 | 22500 | 150 |
| 20x | $512^2$ | 600 | 269 | 1.5 | 1.2 | 18500 | 220 |
| 40x | $256^2$ | 600 | 676 | 1.5 | 2.4 | 31870 | 510 |
| 40x | $512^2$ | 600 | 676 | 0.75 | 1.2 | 48000 | 900 |

***Table S3.1 Comparison of the time needed to scan a square cm of preparation with different lenses.*** *2 FLIM detectors were pooled in one channel, scanning was at the indicated speed, using 0.75 zoom, adjusting the format to ensure identical pixel size (except for the last line in the table). The column "photons/cell" contains an estimate of the number of photons per cell, derived from the data in the other columns, the fact that an average cell covers 250 pixels (at 1.5 µm pixel size), and a 50% reduction to account for dimmer pixels in the nucleus and periphery of the cell. When a single detector is used, photons per cell are halved, whereas other values remain equal. Note that with an increasing number of tiles, a considerable part of the total duration is taken for stage movements.*

### SUPPLEMENTAL MATERIAL 4: DESIGN OF THE LIBRARY

The moderately sized custom-made cAMP gRNA library includes most of the genes that might conceivably be involved in shaping the dynamics of receptor-mediated cAMP signals. The library includes genes that may affect the magnitude of the response, such as G-protein-coupled receptors, $G\alpha_s$, adenylate cyclases, and phosphodiesterases (~40 genes). It also included genes that may alter the longevity of the cAMP signals, for example, genes known or suspected to be involved in receptor internalization, such as receptor kinases, arrestins, clathrin, dynamins, the AP2 complex, and vesicle-trafficking genes (~20 genes). Additionally, we included a number of cAMP effector genes that might function in regulatory feedback loops, as well as genes that might affect the unstimulated baseline cAMP concentration, and a number of controls. Finally, several more distantly related genes were included to increase the potential for discovering new members involved in the pathway. A total of 318 genes were included (See Supplemental Files S4 and S5).

Genes were selected from KEGG[5] and STRING database[6], supplemented by literature reviews[7,8]. Further refinement of the selection was aimed at avoiding essential genes (as these genes are expected to drop out of the screen)[9], and on the expression profiles in HeLa and Hek cells derived from Human Protein Atlas data[10]. In the library description (Supplemental File S4), each gene is linked to key resources like its Hugo Gene Nomenclature Committee (HGNC) ID for comprehensive gene information and its CRISPR screen summary from BioGRID ORCS[11], offering insights into whether it has been identified as a hit in other genome-wide CRISPR screens. Additionally, every gene is connected to its STRING database entry, enabling the exploration of its interaction network and facilitating a deeper understanding of its role within the cAMP pathway. This curated and well-annotated library should support the identification of mostly established and putative regulators of cAMP signaling.
